# Latitudinal cline does not predict variation in the microbiome of wild *Drosophila melanogaster*

**DOI:** 10.1101/2022.02.19.481158

**Authors:** Lucas P. Henry, Julien F. Ayroles

## Abstract

Microbiomes affect many aspects of host biology, but the eco-evolutionary forces that shape their diversity in natural populations remain poorly understood. Geographic gradients, like latitudinal clines, generate predictable patterns in biodiversity at macroecological scales, but whether these macro-scale processes apply to host-microbiome interactions is an open question. To address this question, we sampled the microbiomes of 13 natural populations of *Drosophila melanogaster* along a latitudinal cline in eastern United States. The microbiomes were surprisingly consistent across the cline–latitude did not predict either alpha or beta diversity. Only a narrow taxonomic range of bacteria were present in all microbiomes, indicating that strict taxonomic filtering by the host and neutral ecological dynamics are the primary factors shaping the fly microbiome. Additional temporal sampling across two independent sites revealed significant differences in microbial communities over time, suggesting that local environmental differences that vary at fine spatiotemporal scales are more likely to shape the microbiome. Our findings reveal the complexity of eco-evolutionary interactions shaping microbial variation in *D. melanogaster* and highlight the need for additional sampling of the microbiomes in natural populations along environmental gradients.

## Introduction

The microbiome shapes many aspects of organismal biology, contributing to developmental, physiological, and reproductive traits [1]. However, the evolutionary importance of host-microbe interactions remains enigmatic because the complexities of microbial inheritance and strong ecological context dependence complicate traditional models of evolutionary processes [1–3]. It is also poorly understood how ecological forces that drive patterns of biodiversity at macroscales also apply to host-associated microbiomes [4]. To better understand the evolution of host-microbiome interactions, characterization of the microbiome across natural populations is needed to understand drivers of variation across ecologically relevant conditions.

Clines provide a path forward. Clines are geographic gradients (e.g., latitude or elevation) that vary predictably in abiotic and biotic conditions, resulting in spatially variable selection. One striking pattern is the latitudinal gradient of biodiversity, where species diversity is negatively correlated with latitude [5,6]—whether or not host-associated microbiomes display similar patterns is an open question. A key prediction from the latitudinal gradient suggests that the relative importance of biotic and abiotic factors can vary, where low latitudes with higher diversity lead to more complex biotic interactions that constantly shift fitness optimums [6]. For the microbiome, hosts in low latitudes may select for high microbial diversity to increase metagenomic (host + microbial genomes) diversity to find novel solutions to stressful biotic interactions. At high latitudes, abiotic pressures are more predictable (e.g., cold temperatures), and hosts may select for specific microbes to buffer the abiotic pressures, reducing microbial diversity. For hosts with environmentally acquired microbiomes, the processes that govern microbial acquisition from the environment require special attention. Strict filtering may select for specific microbial species across latitudes, resulting in no relationship between latitude and microbial diversity. Identifying the balance between deterministic and ecologically neutral processes in microbial filtering may also shift over latitude, and sampling both host and environment are essential to understand how microbial diversity varies across a latitudinal cline. Few studies have investigated the latitudinal gradient for host-associated microbes, with the expected negative correlation in leaf fungal endophyte communities across several tree species [7], but inconsistent patterns across several populations of wild house mice [8].

*Drosophila melanogaster* is an excellent model to investigate evolutionary responses in clines. *D. melanogaster* are found from 25ºN to 44ºN along the eastern United States, and populations exhibit both genotypic and phenotypic differentiation along a latitudinal cline [9]. Allele frequency varies across multiple loci like in ethanol detoxification [10] and insulin signaling pathways [11], as well as at the genomic level, with lower diversity at northern latitudes [12]. Phenotypically, fly populations in northern latitudes display variation in life-history traits like increased cold tolerance and enter reproductive diapause to survive the colder climate [9,13,14]. The microbiome may also contribute to differentiation along the cline. The *Drosophila* microbiome is environmentally acquired and relatively simple, with <20 bacteria species mostly from the Acetobacteraceae, Lactobacillales, and Enterobacteriaceae, which affect many aspects of fly physiology [15]. In five populations spanning 4º latitude from the middle to northern end of the cline, *Lactobacillus* increased in northern populations, while *Acetobacter* increased in the south [16]. Furthermore, in experimental mesocosms, inoculating seasonally evolving fly populations with *Lactobacillus* led to an enrichment of alleles associated with northern fly populations [17], suggesting that the microbiome contributes to adaptation across the cline. These studies highlight the potential for the cline to also shape the microbiome in *D. melanogaster*, presenting an excellent opportunity to investigate macro eco-evolutionary forces that shape host-microbiome interactions.

Here, we sampled thirteen natural populations of *D. melanogaster* along 11º latitude in the eastern United States using 16S rRNA amplicon sequencing to characterize the microbiome. Along the cline, we sampled flies, frass, and their rotting fruit substrate at orchards and vineyards. Through sampling at both orchards and vineyards, we can separate how different environments shape microbial variation. By comparing individual flies, frass, and substrate, we can examine how deterministic versus ecologically neutral dynamics in the microbiome vary across the cline. We find that the fly microbiome is dominated by a narrow taxonomic range of bacteria and then subsequently shaped by primarily neutral ecological dynamics. Our results highlight the complexity of environmentally acquired microbiomes and untangling the ecological processes that contribute to microbial variation will be key to understanding host-microbiome evolution.

## Methods

Full details for the materials and methods can be found in the supplement.

### Fly sampling

Flies were collected at orchards and vineyards over three weeks in late September-early October 2018 (Fig. 1). At each site, we collected flies through aspiration over the rotting apples or grapes. The substrate (rotting apples or grapes) was collected into a sterile tube and placed on dry ice. Flies were left in an empty tube for ~two hours to collect the frass. Flies typically egest any food as well as pass transient microbes in the gut over two hours [18]. Flies defecated in the empty vial. After two hours, flies were flipped to a clean vial, and both flies and frass vials were placed on ice for ~4-6 hrs. This comparison enables the differentiation between more stably associated microbes in flies and transient microbes in the frass. Flies were sorted morphologically by species (i.e., discarding non *D. melanogaster* or *simulans*), and frass resuspended in sterile water, then both were frozen on dry ice. When we returned to the lab, all samples were kept at −80ºC until processing.

**Figure 1:**
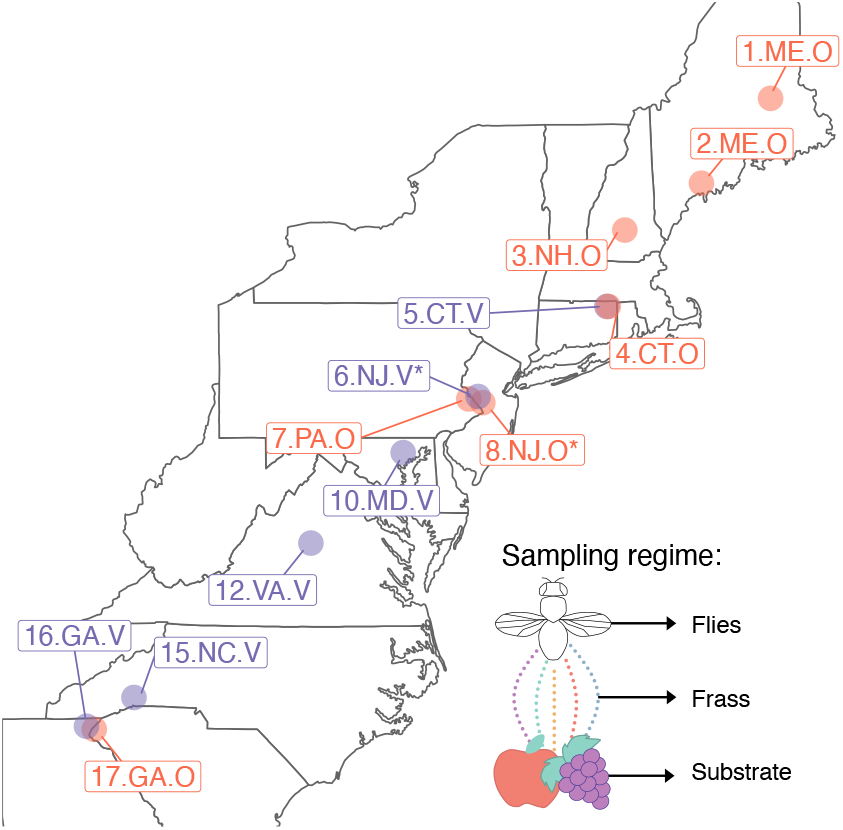
Sampling regime. A) Populations were sampled at each site across 11º latitude over 2 weeks in late September-early October 2018. The stars denote the two sites that were sampled at the end of the growing season (late October). Purple represents vineyards, and red represents apple orchards. The inset diagrams the sampling regime. At each site, we collected flies, frass, and the substrate (either apples or grapes). Individual flies were sequenced, while frass and substrate samples were pooled per site.

DNA was extracted from individual flies using the Zymo Quick-DNA Plus kit, which includes a proteinase K digestion to ensure gram-positive bacteria cells were lysed. The substrate samples yielded low DNA with this extraction kit. We then extracted new DNA for all substrates using the Zymo Quick-DNA fecal/soil microbe kit, as well as a subset of flies to ensure that both DNA extraction methods yielded similar results (Supp. Fig. M1). Microbiomes were characterized through 16S rRNA amplicon sequencing using a two-step dual-indexed approach. Full details can be found in the supplement. In brief, we amplified the V1-V2 region of the 16S rRNA gene (primers in Supp. Table M1), pooled, and then digested with BstZ17l enzyme to deplete Wolbachia amplicons [19]. Libraries were sequenced using 300 bp paired-end reads using the Illumina MiSeq platform at the Princeton University Genomics Core.

Because female *D. melanogaster* and *D. simulans* are morphologically indistinguishable, we performed amplicon sequencing on the COI gene (primers in Supp. Table M1). The proportion of *D. melanogaster* to *D. simulans* varied over the cline (Supp. Fig. M2), and we removed all *D. simulans* from the subsequent analysis. We also removed any site that did not have >10 *D. melanogaster*, resulting in a total of 13 sites with 12-30 flies/site, 1 substrate pool/site, 1 frass pool/site.

### Bioinformatic processing

Frass and substrate samples were sequenced in triplicate, but reads were pooled prior to analysis. Sequences were processed using QIIME2 v.2020.6 [20]. DADA2 [21] was used to cluster into Amplicon Sequence Variants (ASVs), and taxonomy was assigned using the Greengenes reference database [22], trimmed to the 16S rRNA V1-V2 region. Data was imported into phyloseq for visualization and statistical analyses [23]. Sequencing resulted in 3,240,709 reads following quality control, removal of potential contaminants with decontam [24], and filtering out any remaining Wolbachia reads. Samples were rarefied to 500 reads for subsequent analyses (Supp. Fig M3 for rarefaction curves).

### Statistical analyses

For each site, we used latitude, elevation, and the following climatic variables from the nearest weather station [25]: temperature and precipitation from the day, week, month, year, and 5 years preceding the collection (Supp. Table M2). We also collected two measures of land use (human density and home price from [26]) to include in our analyses. Because climate variables are often correlated, we performed principal components analysis on the climate data (Supp. Fig. M4). PC1 (52.7% of variance, Supp. Fig. M4C) was primarily explained by temperature and precipitation, while PC2 (19.6% of variance, Supp. Fig. M4C) was elevation and home price (Supp. Table M3 for PC contributions). PC1 was correlated with latitude (F_2,14_=89.58, p<0.0001, r^2^=0.847, Supp. Fig. M5), and to avoid collinearity with latitude, we only used PC2 in subsequent analyses.

To test for similarity between microbiomes in fly, frass, and substrate, we visualized the microbiomes with principal coordinate analysis (PCoA) using Bray-Curtis dissimilarity from ASVs. We tested significant effects of sample type, latitude, environment PC2, and origin (i.e., orchard or vineyard) on Bray-Curtis dissimilarity using PERMANOVA in vegan [27]. We then grouped ASVs as to whether they were unique to flies or overlapped with frass or substrate for each site. We compared the abundance with membership (i.e., unique or shared across sample types) using a pairwise Wilcox test with p-values adjusted using Benjamini & Hochberg correction for multiple testing.

Next, we tested if alpha and beta-diversity differed across the latitudinal cline. For alpha-diversity, we calculated Shannon diversity and Faith’s phylogenetic diversity on ASVs. For beta-diversity, we calculated Bray-Curtis dissimilarity on each group separately (i.e., only fly or frass or substrate). In general for each of these diversity measures, we fit a statistical model with latitude, origin (orchard/vineyard), environmental PC2, and percent *melanogaster* as fixed effects, with site as random effect. We included percent *melanogaster* in our analyses to control for any potential biases that might be associated with differences in fly species composition. For alpha-diversity, mixed linear models were implemented using lme4 [28]. Faith’s phylogenetic diversity was modeled with gamma distributions with the log link. Beta-diversity was tested using PERMANOVA in vegan [27].

Finally, we performed neutral ecological modeling based on the Sloan model [29]. In brief, the neutral model assumes that all microbes in a regional pool (i.e., across all individuals) are equally able to disperse across individuals. If the prevalence of an ASV in an individual is predicted by the abundance in the regional pool, this is consistent with expectations from neutral dynamics. Microbes above this distribution are associated more frequently with hosts, suggesting positive selection, while microbes below suggest negative interactions or dispersal limitation. The neutral model tests for the fit of prevalence in individuals given a regional pool using non-linear least-squares fitting and was implemented using the tyRa package [30]. We first determined neutrality at the *D. melanogaster* species level by using all fly samples to fit the neutral model. Then, we calculated the neutral fit for each site and tested for significant correlation with latitude and origin through linear regression. ASVs with frequency < 10 were removed prior to modeling.

### Temporal variation

For one vineyard (6.NJ.V) and one orchard (8.NJ.O), we performed additional sampling ~3 weeks later at the end of the October growing season. The later sampling point was compared to the in-season sampling point. Flies, frass, and substrate were collected and stored similarly to the main collections, and sequencing libraries were prepared with the rest of the samples. We tested for the effects of substrate, site, season, and the interaction between site and season for beta diversity (Bray-Curtis dissimilarity) using PERMANOVA [27]. We then calculated neutral fit as above and compared between the early and late season for the orchard and vineyard.

## Results

### Microbiome characterization across the latitudinal cline

Across sites, microbiomes were composed of similar bacteria, but the relative abundance of different genera varied (Fig. 2A). *Gluconobacter* (31.0%), *Commensalibacter* (24.0%), and *Acetobacter* (19.1%) were the most common bacterial genera detected across all sites. These dominant bacterial genera represent a narrow taxonomic range, as all are from the Acetobacteraceae family. Overall, the Acetobacteraceae family comprised 85.1% of bacteria, while the next most abundant family was the Enterobacteriaceae at 4.8% (Supp. Figure R1). Across sample types, flies tended to harbor more genera than either the substrate or frass (Fig. 2A). The frass tended to be enriched for bacteria from an unclassified Pasteurellales genus compared to either flies or the substrate. The substrate microbiome had the least taxonomic richness, with only 1-3 genera detected, primarily from *Acetobacter*, *Gluconoacetobacter*, and *Gluconobacter*.

**Figure 2:**
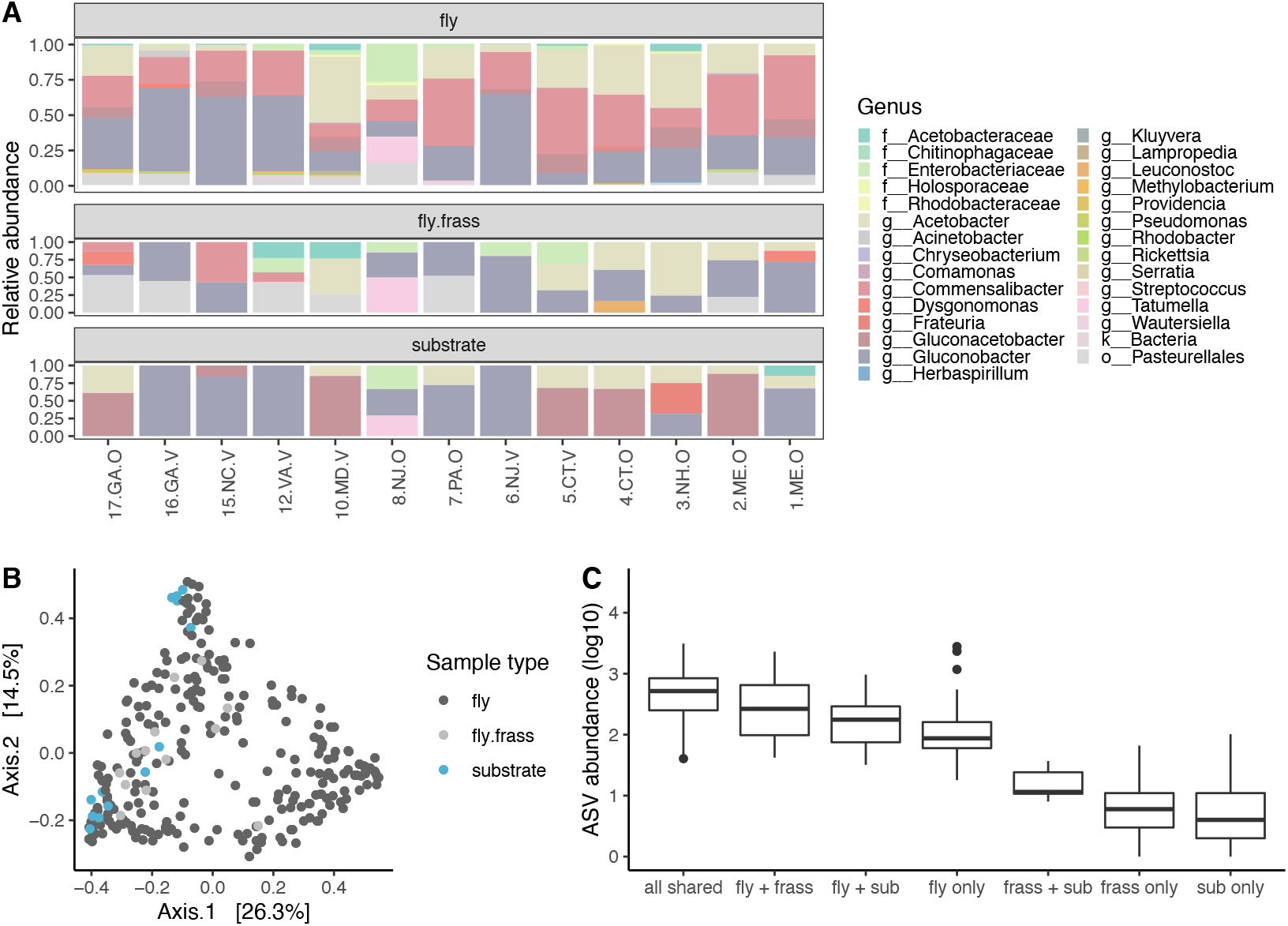
Community composition across sample types along the latitudinal gradient. A) Relative abundance of bacterial genera (>10%). Sites are ordered from south to north. Most genera can be found in multiple sample types, but the relative abundance differed. B) PCoA plot for Bray-Curtis dissimilarity based on ASV composition. For flies, each point represents an individual, while points represent pools of frass or substrate. C) Abundance of ASV overlap between sample types averaged across each site.

Principal coordinates analysis using Bray-Curtis dissimilarity showed substantial overlap between sample types, where frass and substrate cluster among fly samples (Fig. 2B). Sample type explained small, but significant variance between samples (PERMANOVA r^2^ = 0.03, p=0.001, Supp. Table R1). Latitude, climate PC2, and origin (i.e., orchard or vineyard) each explained less than 5% of variance for all microbiomes sampled (Supp. Table R1). A similar trend was observed using Unifrac distance (Supp. Fig. R2, Supp. Table R2).

We examined if ASVs were shared across sample types for each site. If ASVs that are shared are abundant, this would suggest that the environment strongly shapes microbial composition. Conversely, if ASVs that are unique to each group are high abundance, this would suggest stronger filtering, leading to more specific associations between microbes and sample types. ASVs that were shared between all three sample types were the most abundant (Fig. 2C, Table 1). Notably, ASVs unique to flies were more abundant than ASVs shared or unique to frass and substrate. Overall, the results suggest that a combination of environmental determination and filtering may shape the fly microbiome.

**Table 1:** P-values (Benjamani-Hochberg corrected from pairwise Wilcoxon rank sum test) for differences in ASV abundance shared between substrate types.

|  | **all shared** | **fly + sub** | **fly + frass** | **frass + sub** | **fly only** | **frass only** |
| --- | --- | --- | --- | --- | --- | --- |
| fly + sub | 7.90E-10 | - | - | - | - | - |
| fly + frass | 0.01479 | 0.00247 | - | - | - | - |
| frass + sub | 6.30E-06 | 7.90E-06 | 6.30E-06 | - | - | - |
| fly only | 7.10E-16 | 0.00058 | 1.20E-08 | 1.40E-05 | - | - |
| frass only | < 2e-16 | < 2e-16 | < 2e-16 | 0.0035 | < 2e-16 | - |
| sub only | < 2e-16 | < 2e-16 | < 2e-16 | 0.00368 | < 2e-16 | 0.00299 |

### Latitude does not predict alpha or beta diversity in the fly microbiome

We next compared both alpha and beta diversity across the latitudinal gradient in the fly microbiome. Overall, latitude was not correlated with either Faith’s phylogenetic diversity or Shannon diversity (Fig. 3A, Supp. Table R4). Origin and climate PC2 also did not significantly affect alpha diversity measures. While we detected a significant trend for latitude to predict frass phylogenetic diversity (F_2,10_ = 5.60, p=0.02, Supp. Fig. R3), we did not find a significant association between frass and substrate on fly alpha diversity (Supp. Fig. R4). Similarly, for beta diversity, latitude only explained ~1% of variance in the fly microbiome (Fig. 3B, Supp. Table R5). Site explained ~12% of variance, suggesting that fine-scale local conditions shaped the microbiome more than broad-scale, eco-evolutionary interactions across the cline.

**Figure 3:**
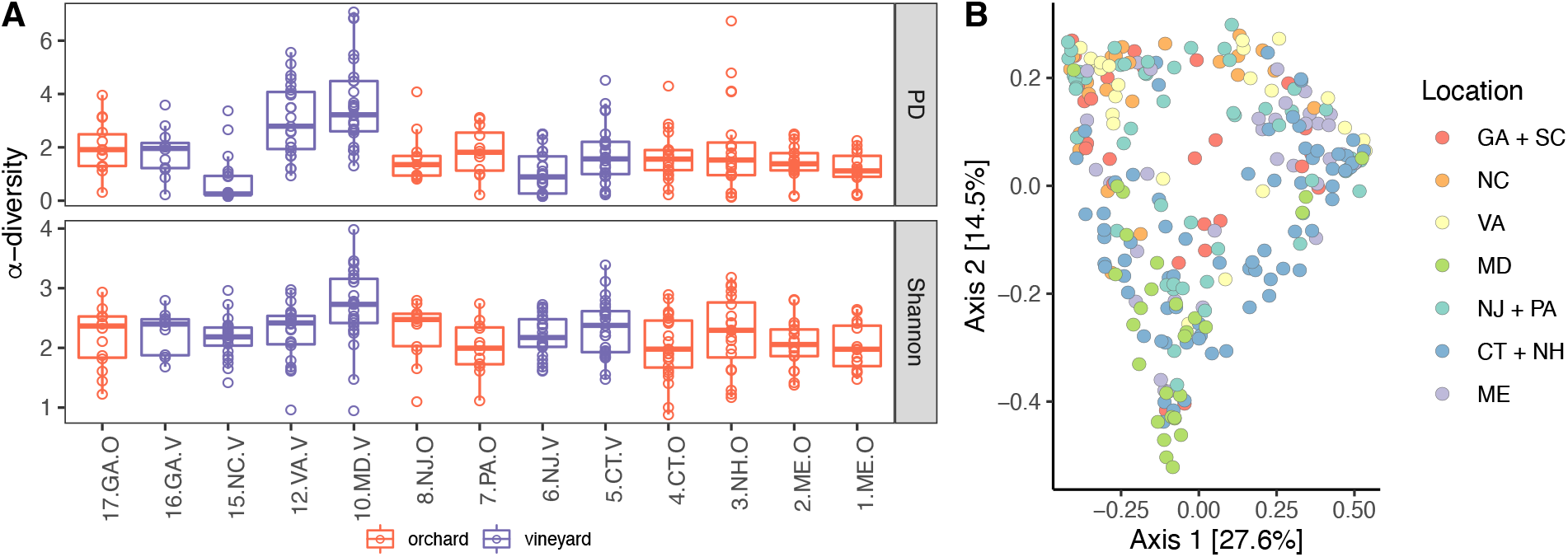
Latitude does not explain variation in alpha or beta diversity. A) Alpha diversity (Shannon and PD= Faith’s phylogenetic diversity) for fly samples from each site. Site is arranged by latitude and colored by orchard/vineyard. B) PCoA plot using Bray-Curtis dissimilarity. Each point represents a fly. Warmer colors represent southern populations, while cooler colors represent north populations (grouped by U.S. state).

### Neutral ecological dynamics dominate the fly microbiome

Because we detected minimal influence of latitude and environmental variables on microbiome composition, we next assessed the balance between deterministic and stochastic processes through neutral ecological modeling. We fit a neutral ecological model [29] that predicts the prevalence of ASVs in individuals given the abundance in a regional pool of microbes from all individuals. ASVs above the prediction are more prevalent than in the regional pool, suggesting positive associations with the host, while those below are less prevalent and suggestive of negative associations.

For all flies across the cline, we find the neutral dynamics dominate the fly microbiome (Fig. 4A, r^2^ = 0.96). The majority of ASVs (93.3%) fit the model’s predictions, while 3.3% were above and 3.4% were below the predicted prevalence. Within each site, the relative importance of neutral dynamics varies, ranging from as low as r^2^ = 0.46 to 0.95 (Fig. 4B). However, neutrality was not predicted by latitude or origin (F_2,10_ = 0.69, p=0.52, Supp. Table R6). Our analysis suggests that the microbiome of *D. melanogaster* is shaped predominantly by neutral dynamics, but local conditions at each site varied in the balance between deterministic and stochastic processes.

**Figure 4:**
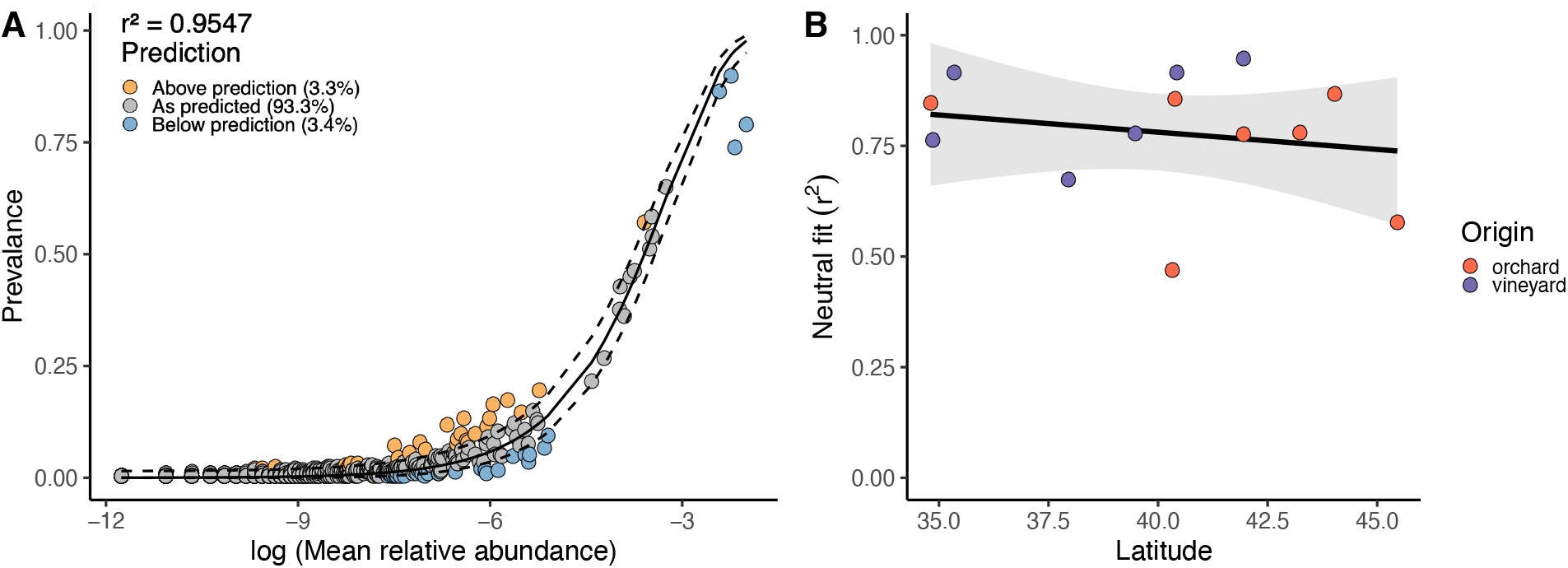
Neutral dynamics dominate the fly microbiome, but vary across each site. A) Fit for neutral community assembly by each ASV across all flies sampled. r^2^ denotes goodness of fit. Point color represents if prevalence in flies was above (red), below (blue), or as predicted (grey) by the neutral model. Dashed lines represent 95% confidence intervals around the model prediction (solid line). B) Neutral model fit (r^2^) across the latitudinal gradient. Each point represents a site, colored by orchard or vineyard.

### Seasonality changes fly microbiome and neutral ecological dynamics

For one vineyard (6.NJ.V) and one orchard (8.NJ.O), we performed additional sampling ~3 weeks later at the end of the October growing season. The microbiome of flies, frass, and substrate shifted over the growing season, but in different ways between the vineyard and orchard (Fig. 5A). For the vineyard, *Commensalibacter* increased in relative abundance from 22.3% to 63.3%, while *Gluconobacter* decreased from 58.1% to 9.2% across all sample types. The opposite occurred in the orchard, where *Gluconobacter* increased from 15.6% to 44.3% across all sample types.

**Figure 5:**
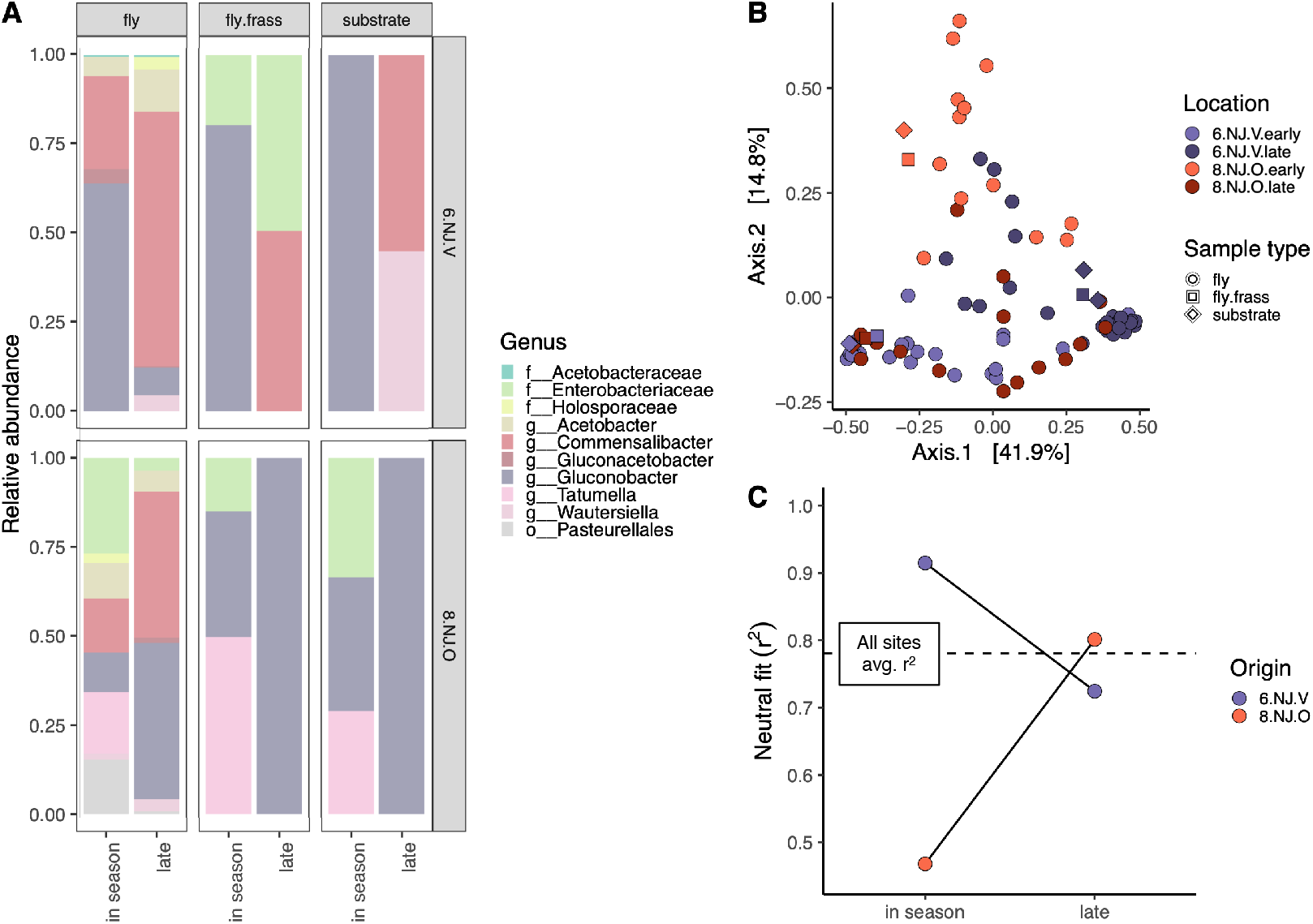
Seasonal dynamics shape fly microbiome at two sites in central New Jersey. A) Relative abundance of genera for the sample types in the vineyard (6.NJ.V) and orchard (8.NJ.O) across two sampling points. B) PCoA plot using Bray-Curtis dissimilarity. Each point represents a fly or pool of frass/substrate. Purple represents samples from the vineyard, while red is for the orchard. Darker colors represent the late season sampling. C) Comparison of neutral fit for the in season and late season sampling across the two sites. The dotted line represents the average neutral fit (r^2^) from all sites sampled in season. complexity of ecological communities could drive microbial dynamics, and more research is necessary to identify how the interactions across ecological scales (i.e., within hosts versus between hosts and species) may vary over clines.

Indeed, the interaction between seasonality and site significantly explained variance in beta diversity between flies (Fig. 5B, Table 2). Site explained 5.1% of variance, seasonality explained 12.7% of variance (PERMANOVA, p=0.001) and the interaction of site x seasonality explained 14.3% of variance (PERMANOVA, p=0.001). Similarly, the contribution of neutral dynamics shifted between the orchard and vineyard over the growing season (Fig. 5C). The orchard had the lowest r^2^ at 0.48 during the earlier sampling period, but increased to 0.80, while the vineyard decreased from 0.92 to 0.78. Intriguingly, for these sites, the neutral fit converged on the average from all sites. Together, this suggests that environmental change can shape the contribution of neutral ecological dynamics that shape variation in the fly microbiome.

**Table 2:** PERMANOVA results for Bray-Curtis dissimilarity in flies during seasonal sampling. Significance codes: 0 ‘***’ 0.001 ‘**’ 0.01 ‘*’ 0.05 ‘.’ 0.1 ‘ ’ 1.

PERMANOVA: Bray-Curtis ~ Substrate + Season + Site + (Season\*Site)
|  | **Df** | **SumsOfSqs** | **MeanSqs** | **F.Model** | **R<sup>2</sup>** | **Pr(>F)** |
| --- | --- | --- | --- | --- | --- | --- |
| Substrate | 2 | 0.7876 | 0.3938 | 2.2194 | 0.03334 | 0.014* |
| Season | 1 | 2.9938 | 2.9938 | 16.8713 | 0.12674 | 0.001*** |
| Site | 1 | 1.2115 | 1.2115 | 6.8271 | 0.05129 | 0.001*** |
| Season:Site | 1 | 3.3679 | 3.3679 | 18.9792 | 0.14258 | 0.001*** |
| Residuals | 86 | 15.2607 | 0.1774 |  | 0.64605 |  |
| Total | 91 | 23.6214 |  |  | 1 |  |

## Discussion

Here, we characterized the microbiome from natural populations of *Drosophila melanogaster* across a latitudinal cline in the eastern United States. Flies harbored bacteria primarily from the Acetobacteraceae family. Latitude did not predict alpha or beta diversity, nor did environmental variables we tested. Importantly, at the two sites we sampled later in the growing season, the microbiome changed significantly and in different ways between the orchard and vineyard. Overall, variation in the fly microbiome was predicted by neutral ecological dynamics. Ecological neutrality, combined with the narrow taxonomic range, suggest that *D. melanogaster* strongly filters for Acetobacteraceae bacteria, but not at finer taxonomic resolution. The composition of bacteria is more likely to depend on the specific microenvironment of the vineyard or orchard. We next discuss the implications for the evolution of host-microbe interactions, particularly when the microbiome is environmentally acquired.

### D. melanogaster harbors specific microbiome across the cline

The microbiomes observed here were primarily composed of bacteria from the Acetobacteraceae family (Fig. 2A, Supp. Fig. R1). Other surveys of wild flies also find Acetobacteraceae as the most common bacteria, including across Europe [31], New York [18], and California [32]. The Acetobacteraceae, like *Gluconobacter*, *Commensalibacter*, and *Acetobacter* species, are all sugar specialists that ferment rotting fruits, like grapes and apples [33]. *Acetobacter* species tend to promote larval development [34] and regulate the conversion of glucose into fat [35], essential for flies living on sugar rich diets like apples and grapes. The dominance of Acetobacteraceae suggests that rotting fruit enriches particular microbes, but at the genus level, different processes affect microbial diversity in flies.

Other studies have suggested *Lactobacillus* should increase at the northern end of the cline. *Lactobacillus* often exerts a suite of effects on fly phenotypes that result in slower life-history traits, which are thought to better match traits with cooler climates [16,17]. Indeed, Walters et al. [16] found that *Lactobacillus* increased in abundance in northern latitudes. However, we only detected *Lactobacillus* at very low abundance, for a total of 0.01% relative abundance across all samples. Notably, *Lactobacillus* was only found at <3% relative abundance in ~10/50 populations across Europe [31]. In kitchens in New York, *Lactobacillus* was only found in 30% of flies sampled with mean relative abundance of 3.6% [18]. Finally, in California, *Lactobacillus* was in only one pool at 0.5% relative abundance [32]. All of these studies found microbiomes dominated by Acetobacteraceae. *Lactobacillus* are also frequently found in laboratory flies, and given their lab prevalence, are thought to form the core microbiome of *D. melanogaster* [15]. Our results highlight how studying wild populations of laboratory model systems in natural environments can reveal surprising insights into the microbiome.

### Alternative drivers of microbial variation across the cline

While latitude and origin (vineyard/orchard) did not explain significant differences in microbial variation (Fig. 3), we do believe we sampled flies in a relevant eco-evolutionary context. First, as discussed above, similar bacteria were found as in other studies of natural fly populations. Second, while not the focus here, we did sample many of the sister species, *D. simulans*, and found significant differences in the proportion of fly species across the cline (Supp. Fig. D1). Higher proportions of *D. melanogaster* were correlated with both higher latitudes and vineyards (r^2^ = 0.48, F_2,14_ = 8.27, p=0.004). The association with vineyards and latitude is not surprising, as increased ethanol tolerance and cold tolerance in *D. melanogaster* over *D. simulans* are well-known examples in ecological differentiation between sister species [36–38]. While we did not find a cline for microbial diversity, we still detected other well-established clinal patterns in *Drosophila*.

We also did not detect strong effects of abiotic variables. Our sampling was performed over 3 weeks to ensure roughly equivalent growing conditions in the orchards and vineyards across the cline. For example, the average temperature for the month of sampling only ranged from 18.5ºC to 23.2ºC across the cline (Supp. Fig. M4). It is possible that we did not detect substantial differences because the environments did not sufficiently differ; additional axes of variation on smaller microenvironmental scales may be more important in shaping microbial diversity. Agricultural practices may be one driver of microenvironmental variation, like insecticides. Insecticides alter the *Drosophila* microbiome through complex changes to nutrition and immunity [39,40]. While we only collected from sites that were actively open to the public (public access is often prohibited immediately following insecticide application), differences in insecticide or other agricultural management practices (e.g., fertilization, fungicides, irrigation) may have influenced microbial diversity.

Biotic interactions among insects may also affect microbial diversity. Rotting apples and grapes are hosts to a wide range of other taxa, as we commonly observed bees, wasps, and ants visiting rotting fruit. While we did not quantitatively assess differences in insect communities, others have noted that *Drosophila* select habitat patches to avoid predators like ants [41]. Furthermore, ecological succession within *Drosophilids* occurs as fruit rots [42]. In the progression of fruit rot, *D. melanogaster* arrives later than *D. simulans* or other species, like *D. hydei*. Notably, ecological succession among flies may be affected by the recent invasion of another Drosophilidae, *Zaprionus indianus*. *Z. indianus* are larger and compete with *D. melanogaster* over oviposition sites and larval food [43]. Competition with *Z. indianus* can alter the evolutionary trajectory of *D. melanogaster* populations during seasonal evolution in field mesocosms [44]. In natural populations, the presence of competition may relegate *D. melanogaster* that cannot compete to less optimal oviposition and food sites, which we observed, but did not systematically quantify and may vary across sites. Differences in the

### Neutrality in environmentally acquired microbiomes

The fly microbiome was surprisingly well predicted by neutral ecological dynamics (r^2^ = 0.96, Fig. 4). Neutrality results from functional equivalence between microbes [29], and in the context of the microbiome, may result from both nonspecific host selection and minimal barriers to microbial dispersal. Microbes that are above the neutral prediction are those that either hosts selectively acquire or are better colonizers, while those below reflect host avoidance or poor colonizers [45,46]. While it is difficult to untangle host selection from microbial colonization, our analyses at two different scales provides some insight into factors that shape neutrality in the microbiome. For all flies, the neutral model closely predicted the dynamics for most ASVs (r^2^=0.96), suggesting that *D. melanogaster* as a species are broadly permissive to colonization by diverse bacteria (though primarily Acetobacteraceae). At each site, the explanatory power of the neutral model ranged from r^2^=0.46-0.95, which suggests local environmental differences generally reduce the contribution of neutral dynamics by either making *Drosophila* more selective or microbes better colonizers. Fly genotype has been shown to shape microbial composition [47], and different microbial genotypes have different propensities to stably colonize flies [48,49]. Additional characterization at finer scale genetic level would provide insight into how interactions between host, microbe, and environment (i.e., G_HOST_ x G_MICRO_ x E) shape the eco-evolutionary dynamics of host-microbe interactions.

The question remains as to why so much of the *Drosophila* microbiome is mostly neutral, as other estimates are often lower from both in *D. melanogaster* from kitchens (r^2^=0.41-0.70 [18]), as well as in zebrafish (r^2^=0.39-0.81 [45]) and sponges (r^2^=0.27-0.66 [46]). The high neutrality likely leads to low fidelity in host-microbe associations. Low fidelity can lead to highly variable associations, which if the host phenotype is responsive, then the microbiome can generate novel phenotypic variation [3]. Theory suggests that this novel phenotypic variation generated from low fidelity associations may also be favored when in variable environments [50]. For *D. melanogaster*, neutrality in the microbiome may enable flies to rapidly adapt to variable environments. Indeed, for the sites that were sampled at the end of the season, the microbiome changed substantially from the initial survey (Fig. 5). Because we only sampled two sites for the late season point, we cannot separate whether this seasonal change reflects stochastic dynamics or more predictable outcomes, like if orchards change differently than vineyards. Our results highlight how organisms that occupy highly variable environments may have flexibility in the microbiome, but more experimental work is necessary to understand the evolutionary benefits from having low fidelity microbial associations.

## Conclusions

Clines provide a powerful framework to investigate how organisms respond to spatially varying patterns of selection, but the microbiome has remained surprisingly understudied. Here, we find that latitude does not predict microbial diversity due to strong neutral ecological dynamics, but the neutrality at sites is temporally variable across the growing season. Characterizing temporal stability in the microbiome through longitudinal sampling across the latitudinal cline is a key research priority. Additionally, we did not investigate the phenotypic effects of microbial variation. Transplanting microbiomes under controlled experimental conditions will show how microbe-induced phenotypic variation may buffer selective pressures that vary along the cline.

To gain mechanistic insight into the many ways the microbiome can shape host biology, we must move beyond correlational studies toward manipulative experiments that inform causal relationships. To that end, several systems like *D. melanogaster*, *C. elegans*, and mice have emerged as amenable models to dissect the effects of the microbiome variation on a wide range of traits in laboratory settings [15]. However, the insight these model systems provide in the laboratory will always be somehow limited. To better understand which ecological and evolutionary forces drive associations between host and microbiome, we should also leverage wild populations. Natural variation in the wild will reveal the fundamental eco-evolutionary processes that drive the complex dynamic governing host-microbiome associations.

## Acknowledgements

We thank the owners of the orchards and vineyards for allowing us to collect samples, A. Bergland and J. Chaston for logistical support, and the Ayroles lab for helpful feedback.

## Data availabilituy

Amplicon sequencing will be uploaded to NCBI [upon acceptance]. Code used to analyze data will be posted on github [upon acceptance].

## SUPPLEMENTARY METHODS

### DNA extraction methods

All samples were initially extracted with the Zymo DNA 96 Plus kit (D4071). Samples were homogenized in 100 μl Solid Tissue Buffer with a single 2.8mm bead (OPS Diagnostics, #089-5000-11) using Talboys High Throughput Homogenizer (#930145). DNA was extracted from individual flies. 1.5 ml of frass samples were centrifuged at 16000 xg for 10 minutes to concentrate, resuspended in 100 μl. Substrate samples were ground using mortar and pestle in liquid nitrogen. 600 μl Solid Tissue Buffer was used for the substrate samples. After 5 minutes of homogenization, the homogenate was digested with proteinase K following for 4-6 hours at 55ºC. After the digest, the frass and substrate samples were aliquoted into three technical replicates, and the DNA extraction continued per the manufacturer’s instructions. We determined DNA concentration using Qubit fluorometric quantification (Thermo Fisher Scientific), and noted that the concentration from the grape samples was often below detection. As we were performing the initial PCR for 16S rRNA V1-V2, we also noted amplification was inconsistent from grapes.

We then used the Zymo Quick-DNA Fecal/Soil Microbiome kit (D6010) on the grapes, 1 apple sample, and four pools of flies (5 females/pool). Flies were included because there were no difficulties in library preparation using the DNA Plus kit and would show if DNA extraction method significantly changed the characterization of the microbiome. The grape and substrate samples were ground in liquid nitrogen using mortar and pestle, and then manufacturer’s instructions were followed. DNA yields were higher and 16S rRNA amplification was more consistent than with the DNA Plus kit. The amplicon libraries were prepared in parallel from both DNA extraction methods and sequenced together on the Illumina MiSeq at the Princeton Genomics Core facilities.

We compared the DNA extracted between the DNA Plus and Fecal/soil kit using PERMANOVA based on Bray-Curtis dissimilarity. For grapes and apples, low yield left very few reads per technical replicate, and so samples were rarefied down to 100 reads. Flies were rarefied to 500 reads. For grapes, at this low coverage, the two extraction methods clustered together (Supp. Fig. M1A), and extraction method did not explain significant variance in beta-diversity (PERMANOVA, F_1,29_ = 0.93, R^2^ = 0.03, p = 0.545). While we only performed one comparison for apples, the two methods cluster together (Supp. Fig. M1B, “North Chester”). For flies, the pooled samples from the Fecal/soil kit clustered within the individual samples from the DNA Plus kit (Supp. Fig. M1C); though we note that there was much more variation among individual flies than within the pools. While extraction method did significantly explain modest variance in beta-diversity (PERMANOVA, F_1,56_ = 6.30, R^2^ = 0.07, p = 0.001), differences between site explained much more variation (PERMANOVA, F_1,56_ = 24.68, R^2^ = 0.28, p = 0.001). DNA extraction methods had modest effects on the characterization of the microbiome, but site differences swamp out technical artefacts.

### COI sequencing

All flies were sequenced for COI to identify between D. melanogaster and D. simulans. Primers can be found in Supp. Table M1. 2 μl DNA (10-40 ng) was used in 10 μl PCR reaction with 5 μl 2X OneTaq HotStart Master Mix (NEB M0484) with 0.4 μl each of the 10 μM forward and reverse primers. Cycling conditions for COI amplification were as follows: 94ºC for 3 min, 25 cycles of 94ºC for 45 s, 55ºC for 1 min, 68ºC for 1 min 30 s, and final extension of 68ºC for 10 min. 2 μl from the first PCR was the template for the second PCR with 5 μl 2X OneTaq HotStart Master Mix and 1μl each of the 5 μM i5 and i7 primers. The cycling conditions for the indexing PCR were: 95ºC for 3 min, 15 cycles of 95ºC for 30 s, 55ºC for 30 s, 68ºC for 30s, and final extension of 68ºC for 5 min.

Subsets of individual flies were checked by gel electrophoresis to confirm amplification along with no amplification in no-template controls. Libraries were pooled by plate with 2 μl/individual and then cleaned using 1X Ampure XP beads. Following bead cleanup, we performed size selection for the correct amplicon size (700 bp) using Zymo Gel DNA Extraction kit (D4007). Libraries were quality checked using Agilent BioAnalyzer and then sequenced on the Illumina MiSeq using 250 bp PE reads.

Sequences were demultiplexed using barcode_splitter [1] and imported into QIIME2 v2020.6 [2]. Sequences were truncated at 240 bp, denoised and ASVs called with the DADA2 plug-in. Species identity was called using a custom reference database consisting of the COI sequences from D. melanogaster and D. simulans. To generate the reference, in silico PCR was performed using the COI PCR primers in the UCSC Genome Browser with D. melanogaster (BDGP Release 6 + ISO1 MT/dm6) and D. simulans (WUGSC mosaic 1.0/droSim1). We confirmed the species assignment of ASVs using BLAST [3].

After taxonomic classification, the data was imported and further analyzed with phyloseq [4]. We removed any ASVs with frequency < 50 across all samples, resulting in 331,845 reads (N=594 flies, avg 558.7 +/− 11.3 SE reads per individual). We obtained a total of 8 ASVs, but most individual flies tended to have just one ASV (Supp. Fig. S1). We only kept flies with >90% relative abundance by a single ASV, which determined the species identity. 17 flies with mixed ASVs (and thus low confidence in species identity) were removed from all subsequent analyses.

### 16S rRNA amplicon library preparation

Flies, frass, and substrate microbiomes were characterized by amplifying the 16S rRNA V1-V2 region. Primers can be found in Supp. Table M1. 1 μl DNA (5-20 ng) was used in 10 μl PCR reaction with 5 μl 2X OneTaq HotStart Master Mix with 0.4 μl of the 10 μM forward and reverse primers. Cycling conditions for 16S rRNA amplification were as follows: 94ºC for 3 min, 25 cycles of 94ºc for 45 s, 50ºC for 1 min, 68ºC for 1 min 30s, and final extension of 68ºC for 10 min. 1 μl from the first 16S PCR reaction was the template for the second PCR to add barcodes (5 μl 2X OneTaq HotStart Master Mix, 1 μl 5 μM i5, 1 μl 5 μM i7 primers). The cycling conditions for the indexing PCR were: 95ºC for 3 min, 15 cycles of 95ºC for 30 s, 55ºC for 30 s, 68ºC for 30s, and final extension of 68ºC for 5 min.

Libraries were pooled by plate with 1.5 μl per individual reaction and then cleaned using 1X Ampure XP beads. Because Wolbachia can be over-represented in sequencing efforts [5] and is not environmentally acquired, we depleted Wolbachia amplicons using BstZ17I restriction digest. The BstZ17I restriction site targets between the V1 and V2 regions to deplete Wolbachia amplicons, is relatively specific to Wolbachia, and does not cleave sites within other commonly found bacteria in the Drosophila microbiome [5]. We confirmed that the BstZ17I restriction digest did not significantly affect the microbiome composition in the substrate samples (PERMANOVA, F_1,16_ = 0.31, r^2^ = 0.02, p=0.91, Supp. Fig. S2). Samples (200-400 ng DNA) were digested with BstZ17I enzyme (NEB R3594) following manufacturer’s instructions (37ºC for 20 min). Digested libraries were separated on 1% agarose gel and then extracted with Zymo Gel DNA Extraction kit. Digested libraries were visualized on Agilent TapeStation and sequenced on the Illumina MiSeq using 250 bp PE reads at the Princeton Genomics Core Facility.

Sequences were demultiplexed using barcode_splitter [1]. The frass and substrate samples were prepared in triplicate, but then were merged for subsequent analyses. All individual flies were sequenced, regardless of species identification. Demultplexed sequences were imported into QIIME2 v2020.6 [2]. Sequences were truncated at 220 bp, denoised and ASVs called with the DADA2 plug-in. Taxonomic classification was performed using the Greengenes reference database [6] trimmed to the 16S rRNA V1-V2 region. Data was then imported into phyloseq [4]. We used the decontam package [7] to remove ASVs associated with no-template controls. We also removed all Wolbachia reads before any statistical analyses were performed.

---

**Supp. Table M1:**
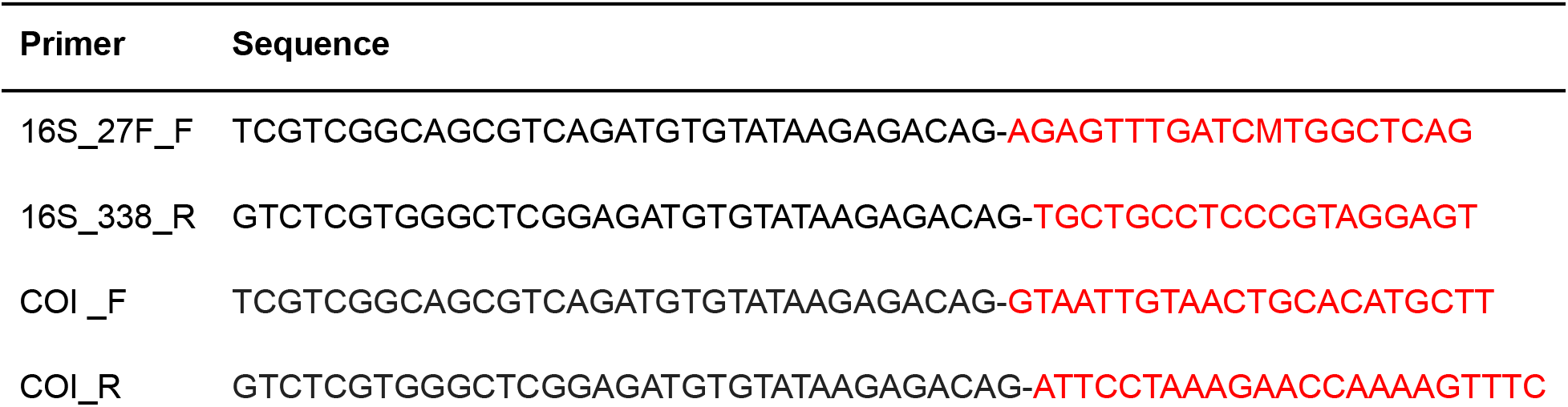
Primers used to generate amplicon sequencing data. The black sequences are the Illumina adapters, while red is locus specific. The 16S rRNA primers are from Simhadri et al. [5], and the COI primers are from Nunes et al. [8].

**Supp. Table M2:**
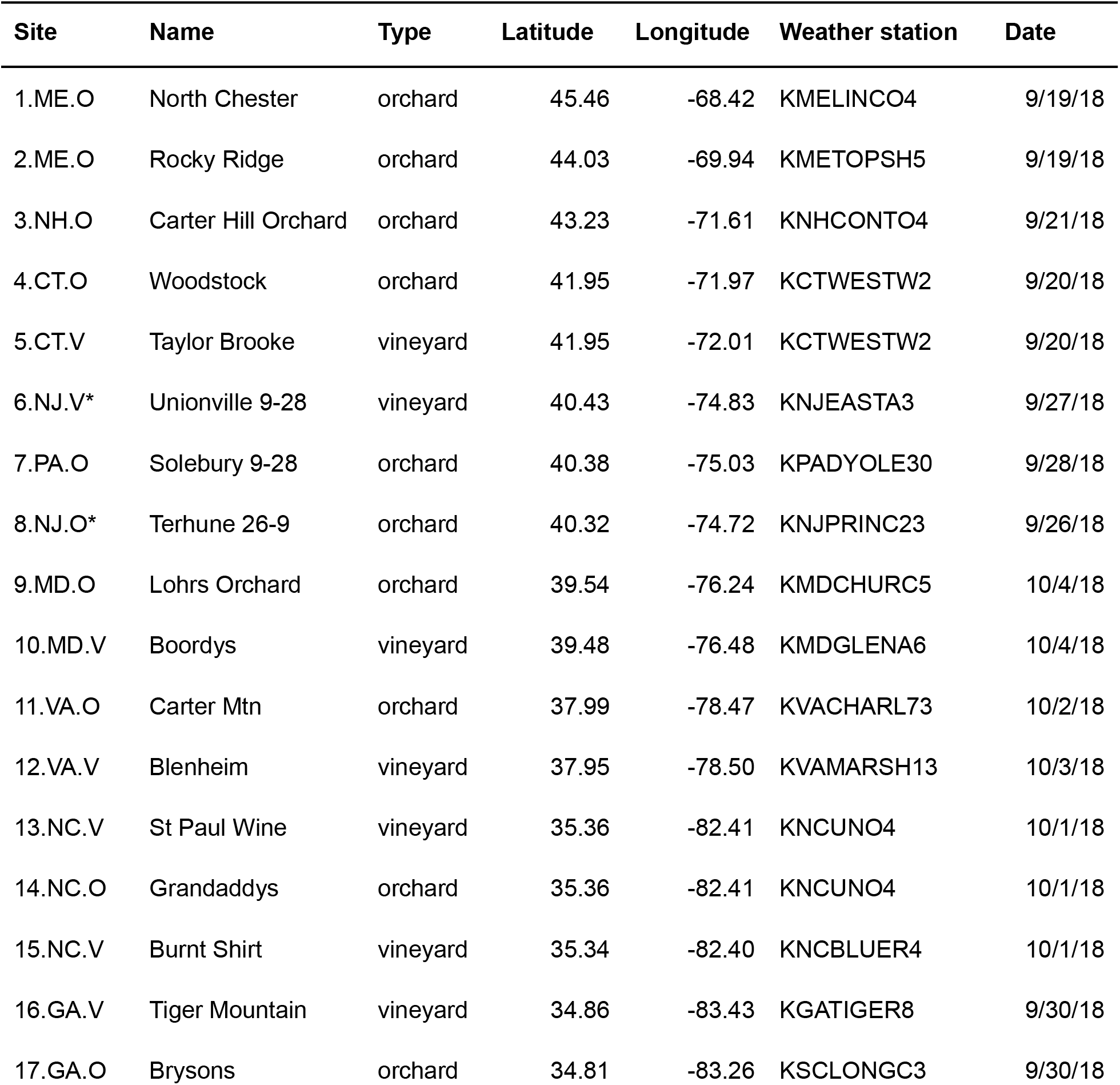
Supp. Table M2: Sampling data for each site. Asterisks denote the two sites that were sampled later in the season.

| Site | Name | Type | Latitude | Longitude | Weather station | Date |
| --- | --- | --- | --- | --- | --- | --- |
| 1.ME.O | North Chester | orchard | 45.46 | -68.42 | KMELINCO4 | 9/19/18 |
| 2.ME.O | Rocky Ridge | orchard | 44.03 | -69.94 | KMETOPSH5 | 9/19/18 |
| 3.NH.O | Carter Hill Orchard | orchard | 43.23 | -71.61 | KNHCONTO4 | 9/21/18 |
| 4.CT.O | Woodstock | orchard | 41.95 | -71.97 | KCTWESTW2 | 9/20/18 |
| 5.CT.V | Taylor Brooke | vineyard | 41.95 | -72.01 | KCTWESTW2 | 9/20/18 |
| 6.NJ.V* | Unionville 9-28 | vineyard | 40.43 | -74.83 | KNJEASTA3 | 9/27/18 |
| 7.PA.O | Solebury 9-28 | orchard | 40.38 | -75.03 | KPADYOLE30 | 9/28/18 |
| 8.NJ.O* | Terhune 26-9 | orchard | 40.32 | -74.72 | KNJPRINC23 | 9/26/18 |
| 9.MD.O | Lohrs Orchard | orchard | 39.54 | -76.24 | KMDCHURC5 | 10/4/18 |
| 10.MD.V | Boordys | vineyard | 39.48 | -76.48 | KMDGLENA6 | 10/4/18 |
| 11.VA.O | Carter Mtn | orchard | 37.99 | -78.47 | KVACHARL73 | 10/2/18 |
| 12.VA.V | Blenheim | vineyard | 37.95 | -78.50 | KVAMARSH13 | 10/3/18 |
| 13.NC.V | St Paul Wine | vineyard | 35.36 | -82.41 | KNCUNO4 | 10/1/18 |
| 14.NC.O | Grandaddys | orchard | 35.36 | -82.41 | KNCUNO4 | 10/1/18 |
| 15.NC.V | Burnt Shirt | vineyard | 35.34 | -82.40 | KNCBLUER4 | 10/1/18 |
| 16.GA.V | Tiger Mountain | vineyard | 34.86 | -83.43 | KGATIGER8 | 9/30/18 |
| 17.GA.O | Brysons | orchard | 34.81 | -83.26 | KSCLONGC3 | 9/30/18 |

**Supp. Table M3:** Contributions (in percentage) of climate variables to PC1 and PC2.

| Climate variable | PC1 | PC2 |
| --- | --- | --- |
| elevation | 3.85 | 15.15 |
| zip.density | 1.06 | 5.76 |
| zip.homeprice | 0.15 | 22.09 |
| day.high | 6.70 | 5.37 |
| day.avg | 8.62 | 2.23 |
| day.low | 7.42 | 0.08 |
| day.range | 1.24 | 10.64 |
| month.avg.high | 1.00 | 1.06 |
| month.avg.avg | 8.65 | 0.14 |
| month.avg.low | 9.88 | 0.11 |
| month.avg.range | 6.71 | 0.72 |
| temp.month | 10.67 | 0.03 |
| precip.month | 0.09 | 19.88 |
| temp.yr | 10.72 | 0.02 |
| precip.yr | 7.02 | 5.06 |
| temp.5yr | 10.75 | 0.05 |
| precip.5yr | 5.48 | 11.62 |

**Supp. Table R1:** PERMANOVA results for Bray-Curtis dissimilarity between sample types, latitude, origin, and climate PC2. Significance codes: 0 ‘***’ 0.001 ‘**’ 0.01 ‘*’ 0.05 ‘.’ 0.1 ‘ ’ 1

PERMANOVA: Bray-Curtis ~ Substrate + Latitude + Origin + Climate PC2 + (strata=site)
|  | Df | SumsOfSqs | MeanSqs | F.Model | R2 | Pr(>F) |
| --- | --- | --- | --- | --- | --- | --- |
| Substrate | 2 | 2.527 | 1.26341 | 4.7571 | 0.03087 | 0.001 *** |
| Latitude | 1 | 2.589 | 2.58883 | 9.7477 | 0.03162 | 0.001 *** |
| Origin | 1 | 1.451 | 1.45133 | 5.4647 | 0.01773 | 0.001 *** |
| Climate PC2 | 1 | 1.999 | 1.99866 | 7.5256 | 0.02441 | 0.001 *** |
| Residuals | 276 | 73.301 | 0.26558 |  | 0.89537 |  |
| Total | 281 | 81.867 |  |  | 1 |  |

**Supp. Table R2:**
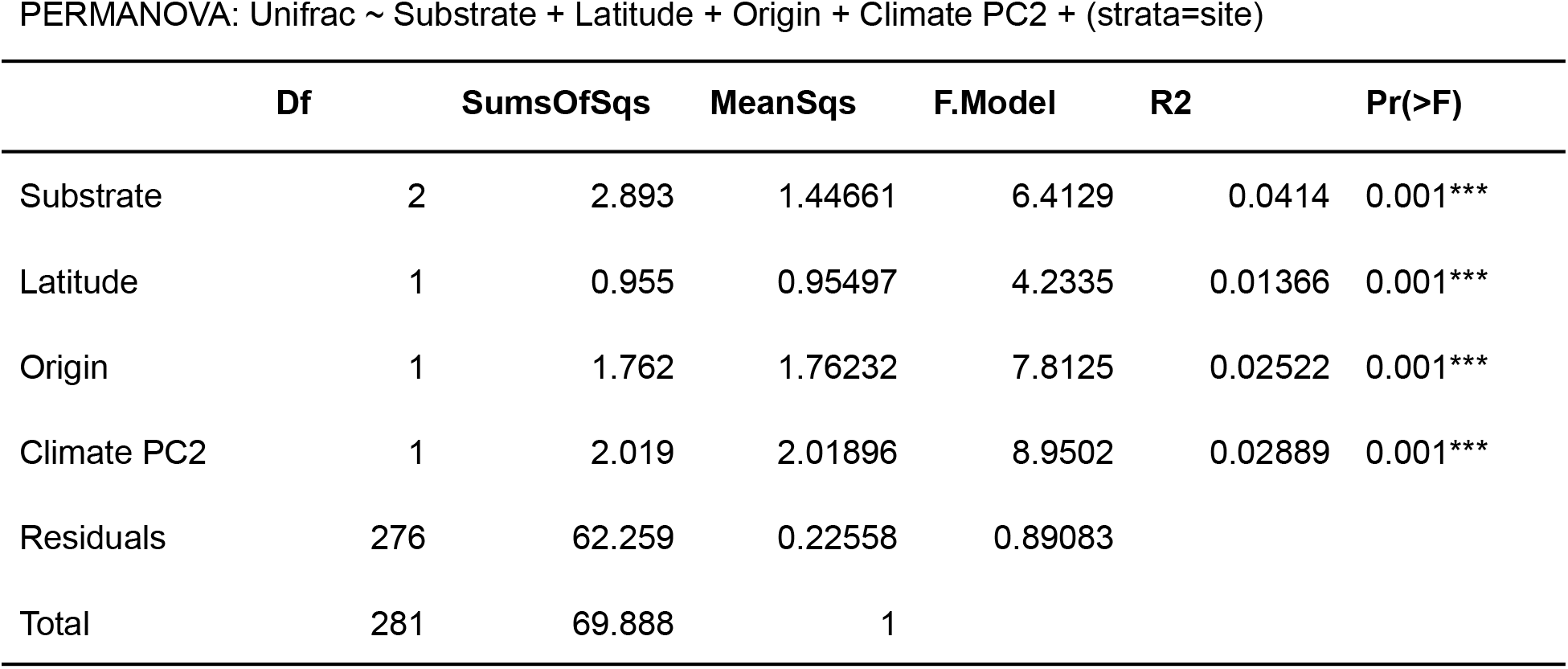
PERMANOVA results for Unifrac between sample types, latitude, origin, and climate PC2. Significance codes: 0 ‘***’ 0.001 ‘**’ 0.01 ‘*’ 0.05 ‘.’ 0.1 ‘ ’ 1

PERMANOVA: Unifrac ~ Substrate + Latitude + Origin + Climate PC2 + (strata=site)
|  | Df | SumsOfSqs | MeanSqs | F.Model | R2 | Pr(>F) |
| --- | --- | --- | --- | --- | --- | --- |
| Substrate | 2 | 2.893 | 1.44661 | 6.4129 | 0.0414 | 0.001*** |
| Latitude | 1 | 0.955 | 0.95497 | 4.2335 | 0.01366 | 0.001*** |
| Origin | 1 | 1.762 | 1.76232 | 7.8125 | 0.02522 | 0.001*** |
| Climate PC2 | 1 | 2.019 | 2.01896 | 8.9502 | 0.02889 | 0.001*** |
| Residuals | 276 | 62.259 | 0.22558 | 0.89083 |  |  |
| Total | 281 | 69.888 | 1 |  |  |  |

**Supp. Table R3:** P-values (Benjamani-Hochberg corrected from pairwise Wilcoxon rank sum test) for differences in ASV abundance shared between substrate types. **currently table 1**

|  | all shared | fly + sub | fly + frass | frass + sub | fly only | frass only |
| --- | --- | --- | --- | --- | --- | --- |
| fly + sub | 7.90E-10 | - | - | - | - | - |
| fly + frass | 0.01479 | 0.00247 | - | - | - | - |
| frass + sub | 6.30E-06 | 7.90E-06 | 6.30E-06 | - | - | - |
| fly only | 7.10E-16 | 0.00058 | 1.20E-08 | 1.40E-05 | - | - |
| frass only | < 2e-16 | < 2e-16 | < 2e-16 | 0.0035 | < 2e-16 | - |
| sub only | < 2e-16 | < 2e-16 | < 2e-16 | 0.00368 | < 2e-16 | 0.00299 |

**Supp. Table R4:** Fixed effects for alpha-diversity measures in flies. Faith’s phylogenetic diversity was modeled using the gamma distribution with log link, while Shannon diversity was modeled with normally distributed residuals. Significance codes: 0 ‘***’ 0.001 ‘**’ 0.01 ‘*’ 0.05 ‘.’ 0.1 ‘ ’ 1

|  | Estimate | Std. Error | t value | Pr(>\|z\|) |
| --- | --- | --- | --- | --- |
| (Intercept) | 1.39597 | 1.77112 | 0.788 | 0.4306 |
| Latitude | -0.01684 | 0.04715 | -0.357 | 0.7209 |
| Origin | 0.0264 | 0.37498 | 0.07 | 0.9439 |
| Climate PC2 | -0.11731 | 0.06872 | -1.707 | 0.0878 . |
| % melanogaster | -0.3053 | 0.78164 | -0.391 | 0.6961 |

Formula: Shannon ~ Latitude + Origin + Climate PC2 + % melanogaster + (1|site)
|  | Estimate | Std. Error | df | t value | Pr(>\|t\|) |
| --- | --- | --- | --- | --- | --- |
| (Intercept) | 2.450027 | 0.628518 | 17.856768 | 3.898 | 0.00107 ** |
| Latitude | -0.008364 | 0.016597 | 17.134022 | -0.504 | 0.62073 |
| Origin | 0.156949 | 0.132146 | 17.051964 | 1.188 | 0.25124 |
| Climate PC2 | -0.045636 | 0.02473 | 18.693235 | -1.845 | 0.08089 . |
| % melanogaster | 0.035666 | 0.282474 | 19.368932 | 0.126 | 0.90083 |

**Supp. Table R5:** PERMANOVA results for Bray-Curtis dissimilarity in flies. Significance codes: 0 ‘***’ 0.001 ‘**’ 0.01 ‘*’ 0.05 ‘.’ 0.1 ‘ ’ 1

PERMANOVA: Bray-Curtis ~ Latitude + Origin + Climate PC2 + % melanogaster + Site
|  | Df | SumsOfSqs | MeanSqs | F.Model | R2 | Pr(>F) |
| --- | --- | --- | --- | --- | --- | --- |
| Latitude | 1 | 0.665 | 0.66475 | 2.6365 | 0.00918 | 0.007** |
| Origin | 1 | 0.209 | 0.20884 | 0.8283 | 0.00288 | 0.567 |
| Climate PC2 | 1 | 0.269 | 0.26911 | 1.0674 | 0.00371 | 0.322 |
| % melanogaster | 1 | 1.178 | 1.17848 | 4.6741 | 0.01627 | 0.002** |
| Site | 8 | 8.857 | 1.10714 | 4.3912 | 0.12226 | 0.001*** |
| Residuals | 243 | 61.268 | 0.25213 |  | 0.8457 |  |
| Total | 255 | 72.446 |  |  | 1 |  |

**Supp. Table R6:** Summary statistics for linear regression between the neutral fit (r^2^) and latitude and origin.

|  | Estimate | Std. Error | t value | Pr(>\|t\|) |
| --- | --- | --- | --- | --- |
| (Intercept) | 0.804901 | 0.560402 | 1.436 | 0.181 |
| Latitude | -0.001617 | 0.013456 | -0.12 | 0.907 |
| Origin | 0.087992 | 0.089832 | 0.98 | 0.35 |

**Supp. Table R7:**
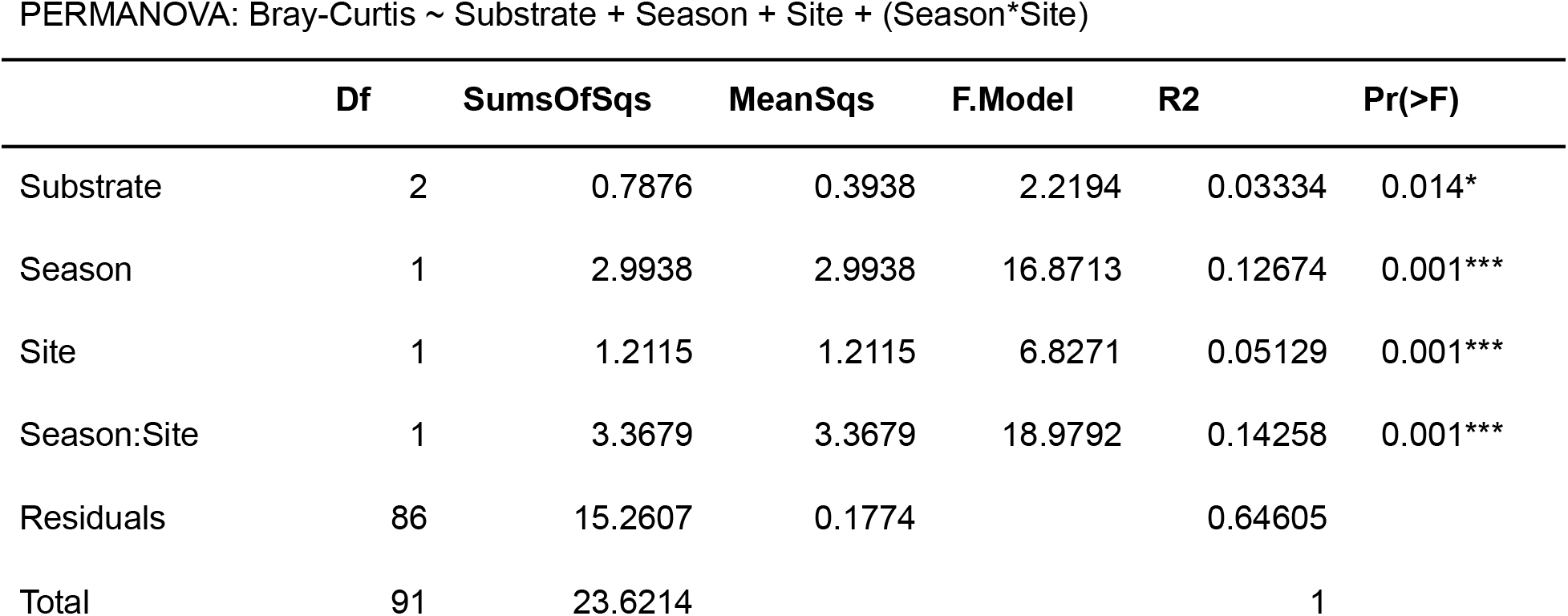
PERMANOVA results for Bray-Curtis dissimilarity in flies during seasonal sampling. Significance codes: 0 ‘***’ 0.001 ‘**’ 0.01 ‘*’ 0.05 ‘.’ 0.1 ‘ ’ 1. **currently table 2**

**Supp. Fig. M1:**
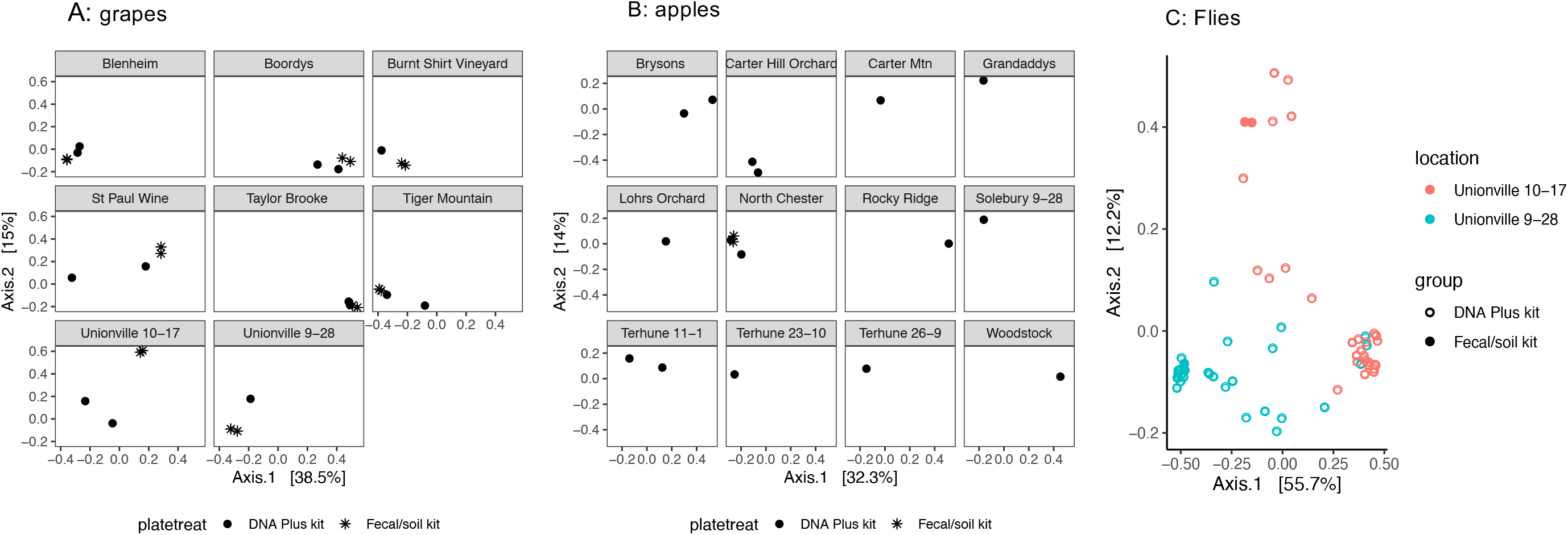
Effects of DNA extraction kit on differences between microbiomes. Plots show PCoA plots based on Bray-Curtis dissimilarity. A) For grapes, there was no significant difference between the extraction kits (PERMANOVA, F_1,29_ = 0.93, R^2^ = 0.03, p = 0.545). Plots are faceted by vineyard name. B) Apples did not exhibit the same difficulty from the DNA Plus extraction kit. Only one location (North Chester) was extracted with the Fecal/soil kit, and the point clusters with the other samples. C) For flies, we only tested on pools of flies using the Fecal/Soil kit, while we did individuals using the DNA Plus kit. Data is shown for the same site at two different sampling points. The pools still fall within samples that match the same time point, though there is more variation among individual fly sequences. The effects of DNA extraction were modest (PERMANOVA, F_1,56_ = 6.30, R^2^ = 0.07, p = 0.001) compared to site differences (PERMANOVA, F_1,56_ = 24.68, R^2^ = 0.28, p = 0.001).

**Supp. Fig. M2:**
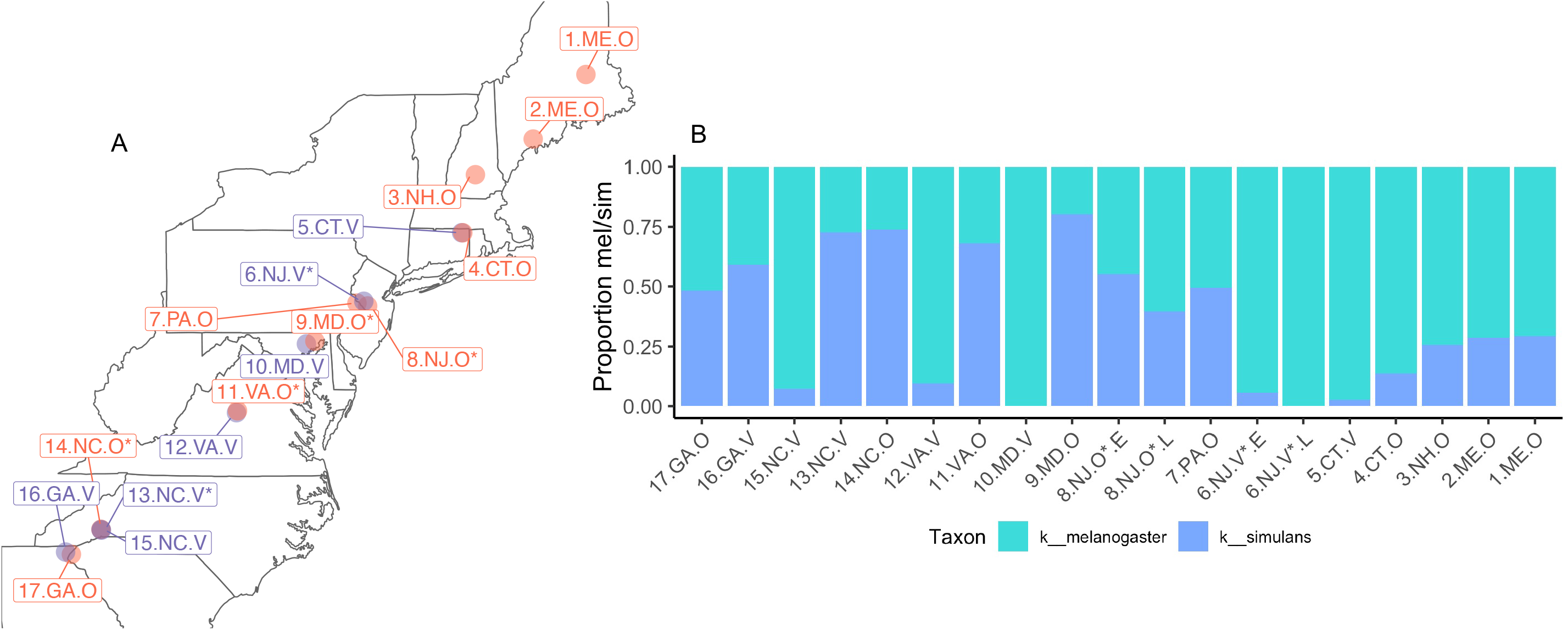
Map of all sites sequenced with the proportion of D. melanogaster and D. simulans across the cline. A) This map shows all the sites sampled. Points are colored by orchard (red) or vineyard (purple). We note that four sites (9.MD.O, 11.VA.O, 13.NC.V, 14.NC.O) had fewer than 10 D. melanogaster and were removed from the analyses in the main text. B) Proportion of D. melanogaster and D. simulans across the cline. All 17 sites are displayed and ordered by latitude.

**Supp. Fig. M3:**
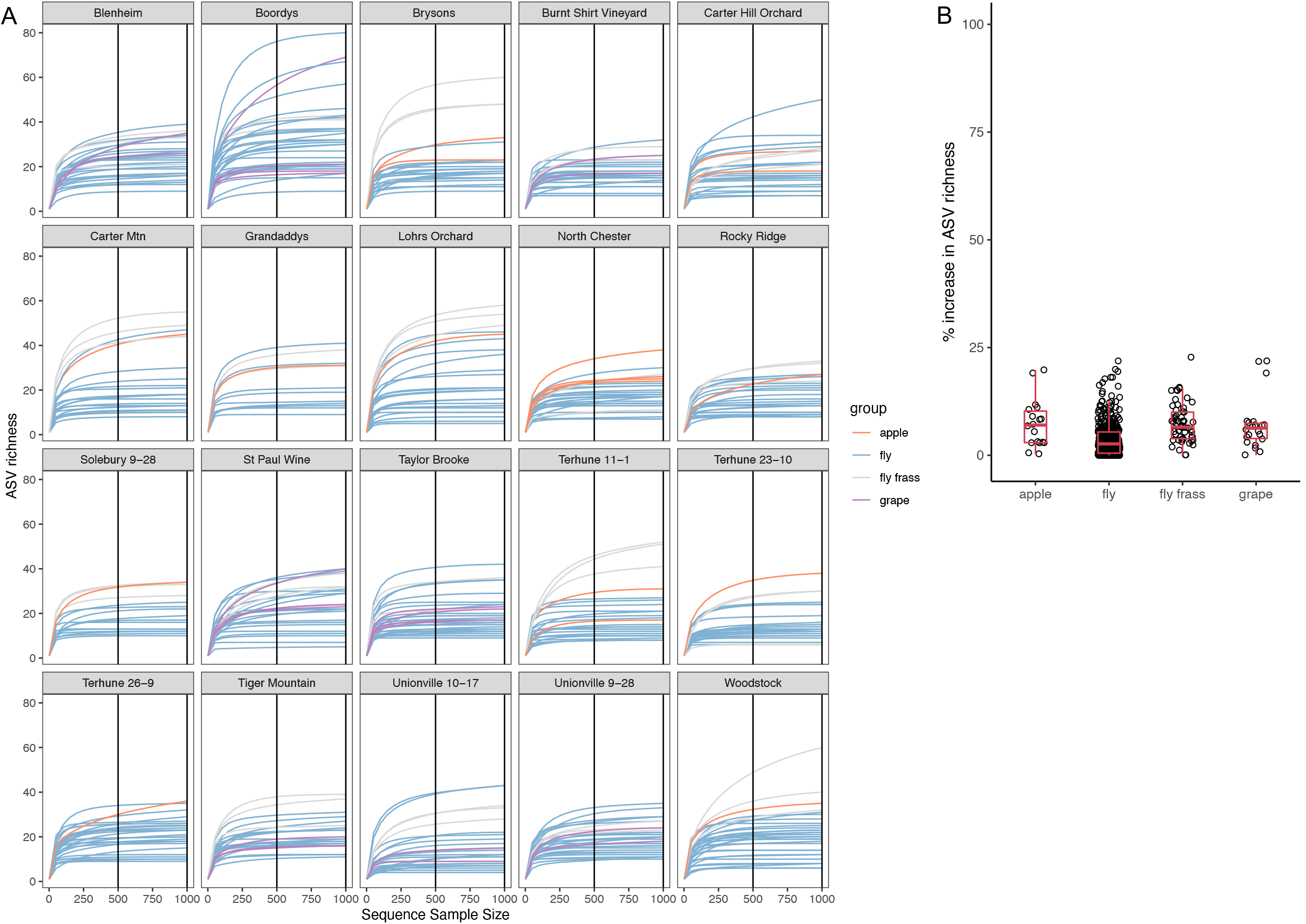
Rarefaction analysis. A) Rarefaction plots shown by site. Each line represents an individual fly (blue), pool of frass (gray), apple (red), or grape (purple). Black vertical lines show 500 or 1000 reads. Plots are faceted by orchard/vineyard for ease of visualization. In general, especially for flies, discovery of new ASVs plateaus at <250 reads/sample. B) Percent increase in ASV richness between rarefying at 500 and 1000 reads. Each point represents an individual, while the red shows boxplots. While in general rarefying to 500 reads/sample is sufficient to capture most of the ASVs, there are some cases, especially for substrates, that the ASV richness did not plateau. At most, increasing rarefaction would increase the ASVs by 25%, but this would have minimal impacts on our interpretation of microbiomes, especially for flies.

**Supp. Fig. M4:**
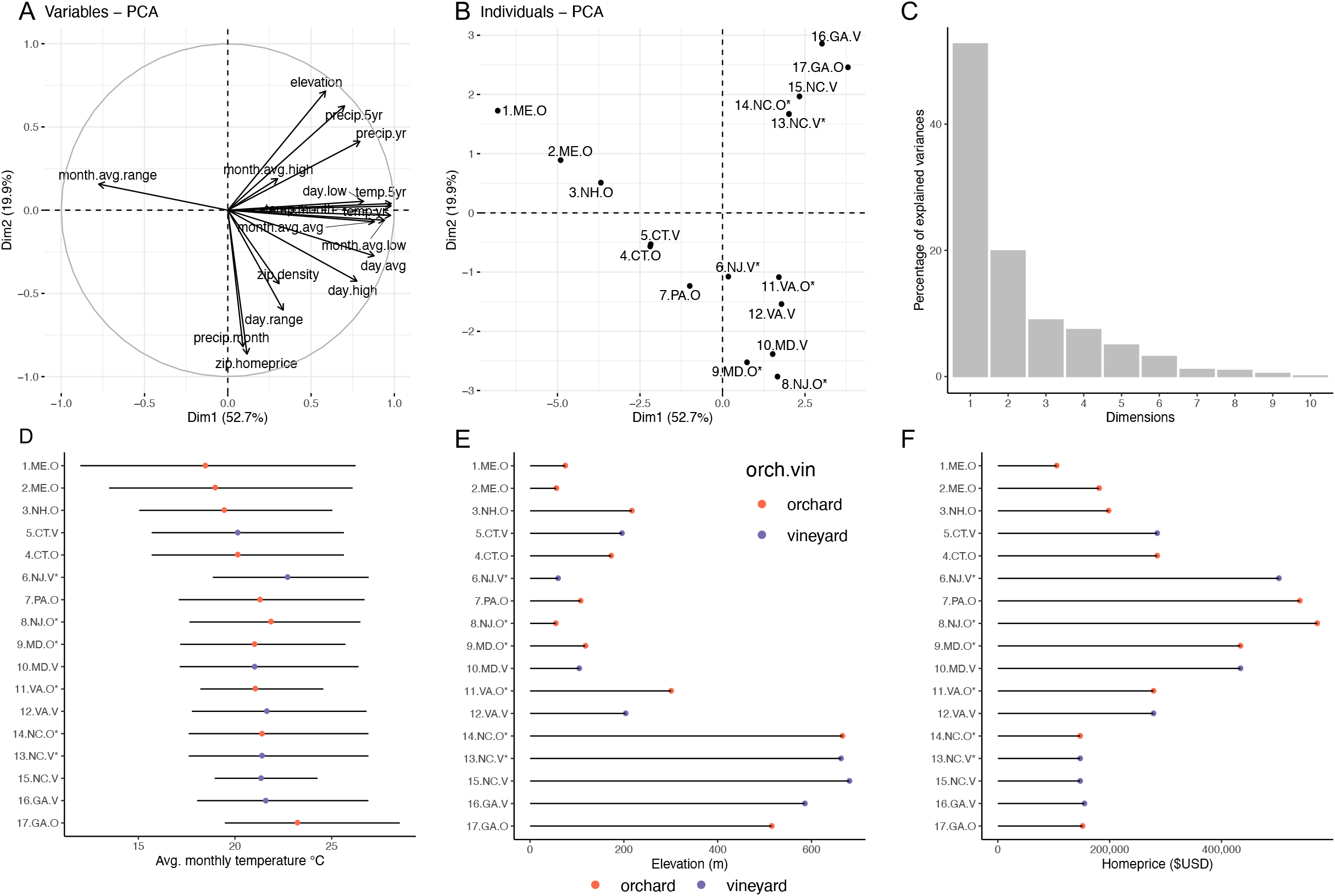
Climatic variables across the cline. A) Principal component analysis of climatic variables. PC1 is primarily temperature and precipitation, while PC2 is predominantly elevation and homeprice. B) PC plot showing each site. C) Scree plot for the top 10 axes. Most variation in the environment is explained by PC1 (52.7%) and then PC2 (19.9%). Plots D, E, F show the range for some in environmental parameters. Points are colored by orchard (red) or vineyard (purple). Sites are arranged by latitude (top=north, bottom=south). D) Monthly temperature range for all locations sampled along the cline. Point represents the average monthly temperature and line shows the range in low and high temperatures for the month before sampling. E) Elevation for each site. F) Average home price by zip code in which the site was located.

**Supp. Fig. M5:**
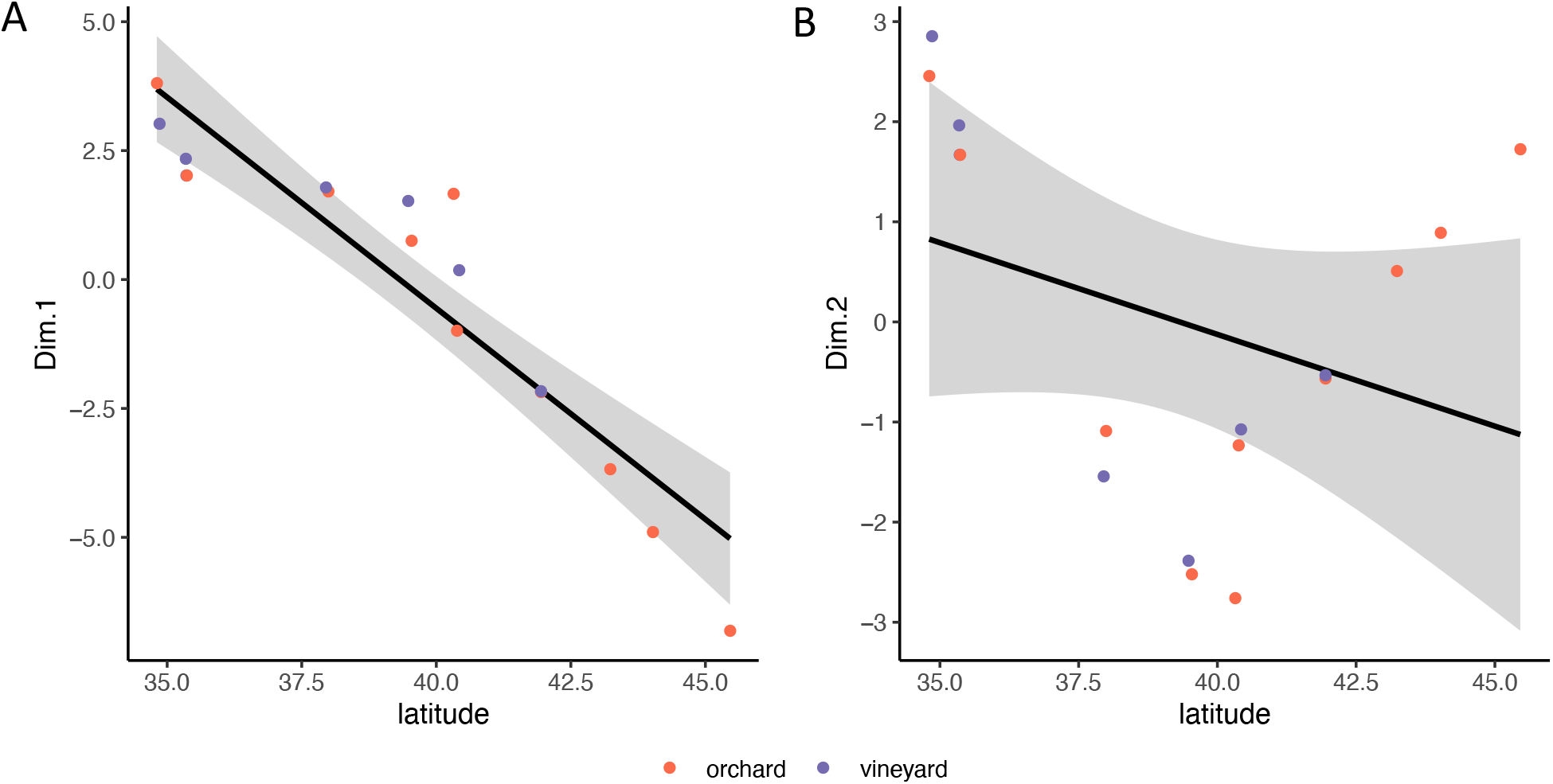
Correlation between climate PCs and latitude. A) PC1 (labeled Dim. 1 here) is correlated with latitude (model: PC1 ~ latitude + origin, F_2,14_=89.58, p<0.0001, r^2^=0.847). Origin (orchard vs vineyard) was not significantly associated with variance in Dim. 1. B) PC2 (labeled Dim. 2 here) was not significantly correlated with latitude (model: PC2 ~ latitude + origin, F_2,14_=0.9337, p=0.4163, r^2^=-0.008).

**Supp. Fig. R1:**
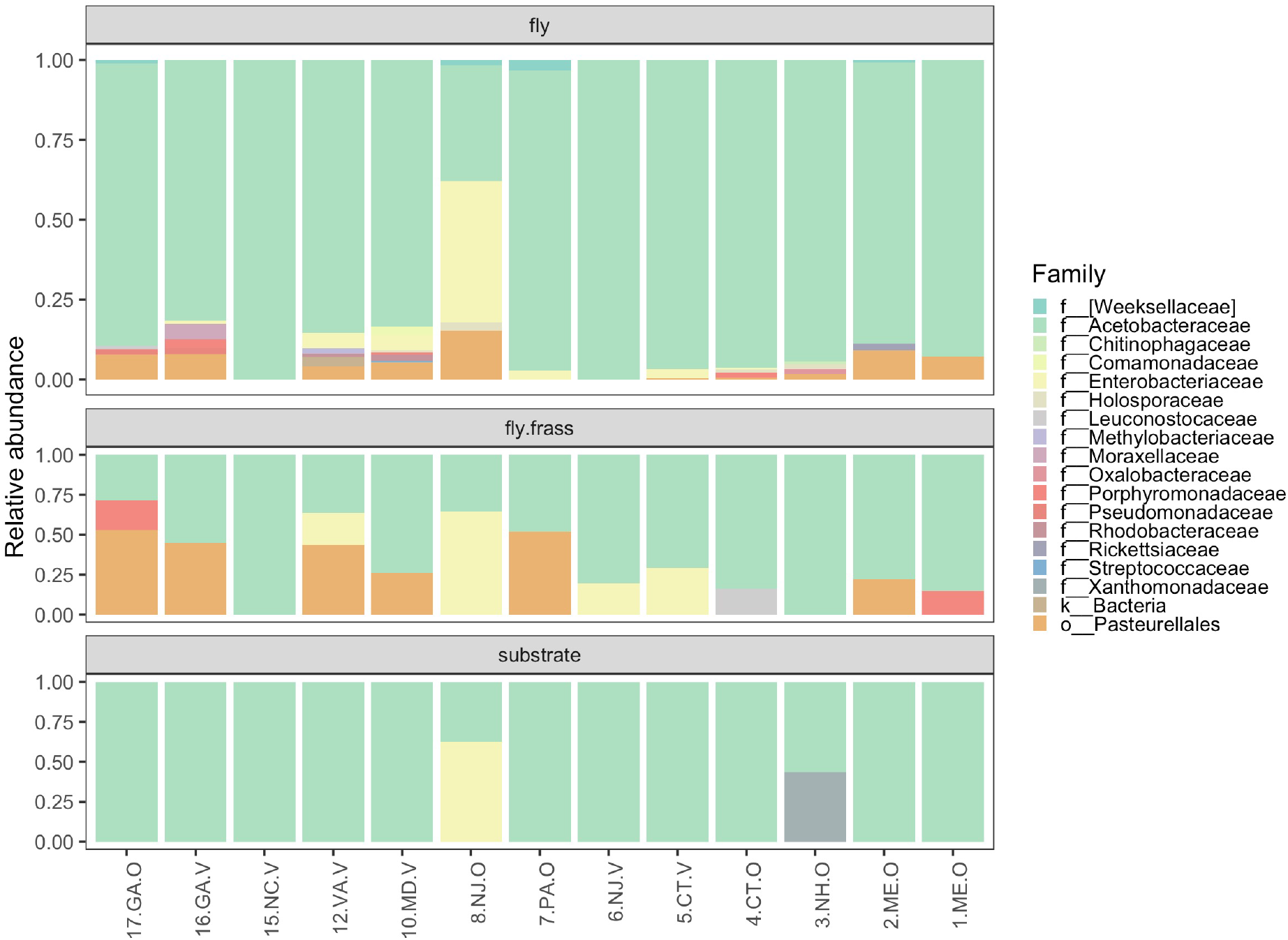
Microbiome composition at the family level. Sites are ordered from south to north. Acetobacteraceae comprised ~85% of all bacteria across all sample types.

**Supp. Fig. R2:**
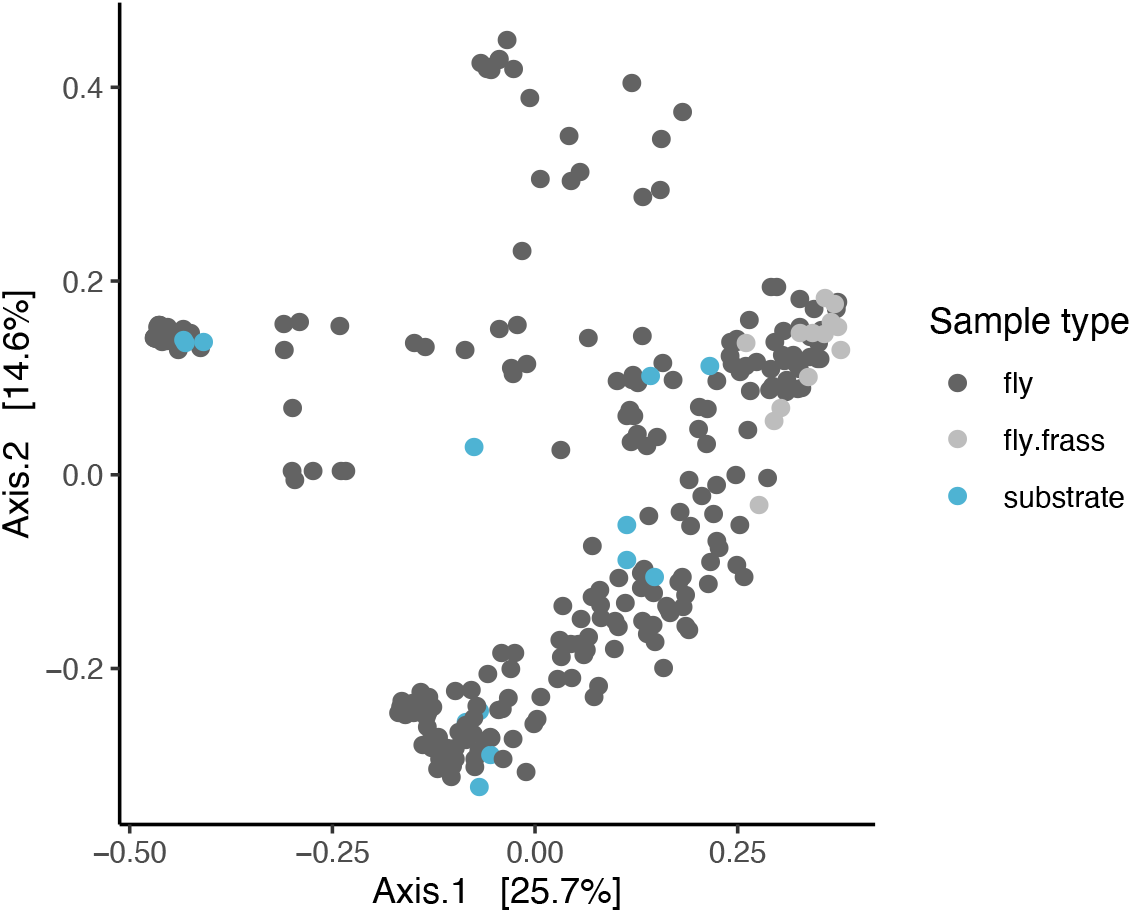
PCoA plot using Unifrac distance. For flies, each point represents an individual while frass and substrate are pools per site. Sample type explained modest by significant amounts of variance between samples (PERMANOVA, F_2,276_ = 6.412, r^2^ = 0.041, p=0.001). Similarly, latitude (, r^2^ = 0.014), origin (r^2^ = 0.025), and climate PC2 (r^2^ = 0.029) explained significant, but low amounts of variance in beta-diversity. These results qualitatively match the findings from Bray-Curtis dissimilarity.

**Supp. Fig. R3:**
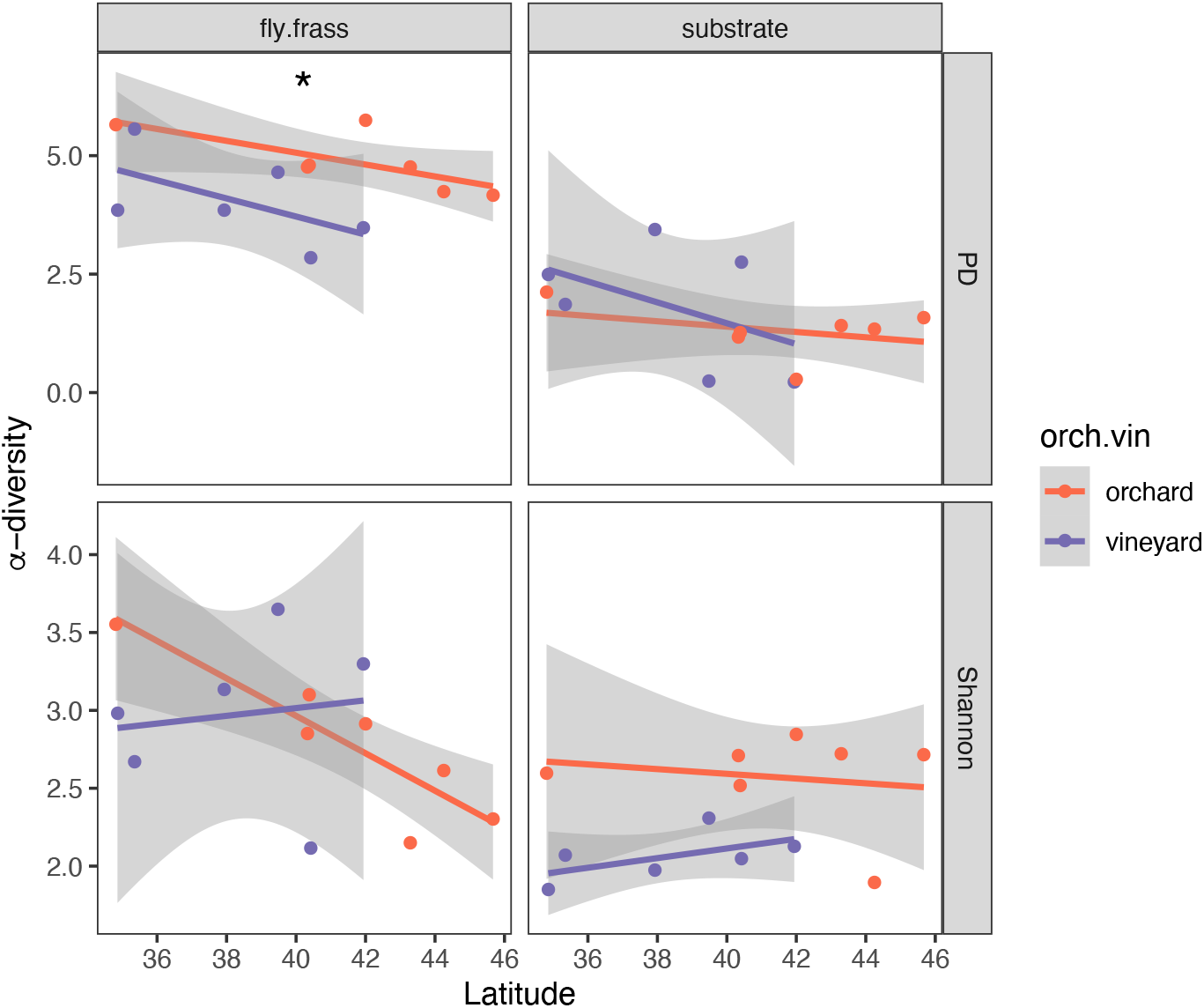
Correlations in alpha diversity between latitude and frass or substrate. Each point represents a pool of frass or substrate from each site, colored by orchard (red) or vineyard (purple). Each comparison was independent and was modeled as: alpha-diversity measure ~ latitude + origin (orchard/vineyard). The asterisk denotes the only significant correlation between phylogenetic diversity (PD) in the fly frass and latitude (F_2,10_ = 5.599, r^2^ = 0.4339, p=0.02337). Shannon diversity was not correlated with latitude in the fly frass (F_2,10_ = 1.551, r^2^ = 0.08, p=0.259). For the substrate, there was no relationship for phylogenetic diversity (F_2,10_ = 1.237, r^2^ = 0.043, p=0.3217). While there was no relationship with latitude for Shannon diversity in the substrate, orchards did have significantly higher diversity than vineyards (t=-2.993, p=0.0135).

**Supp. Fig. R4:**
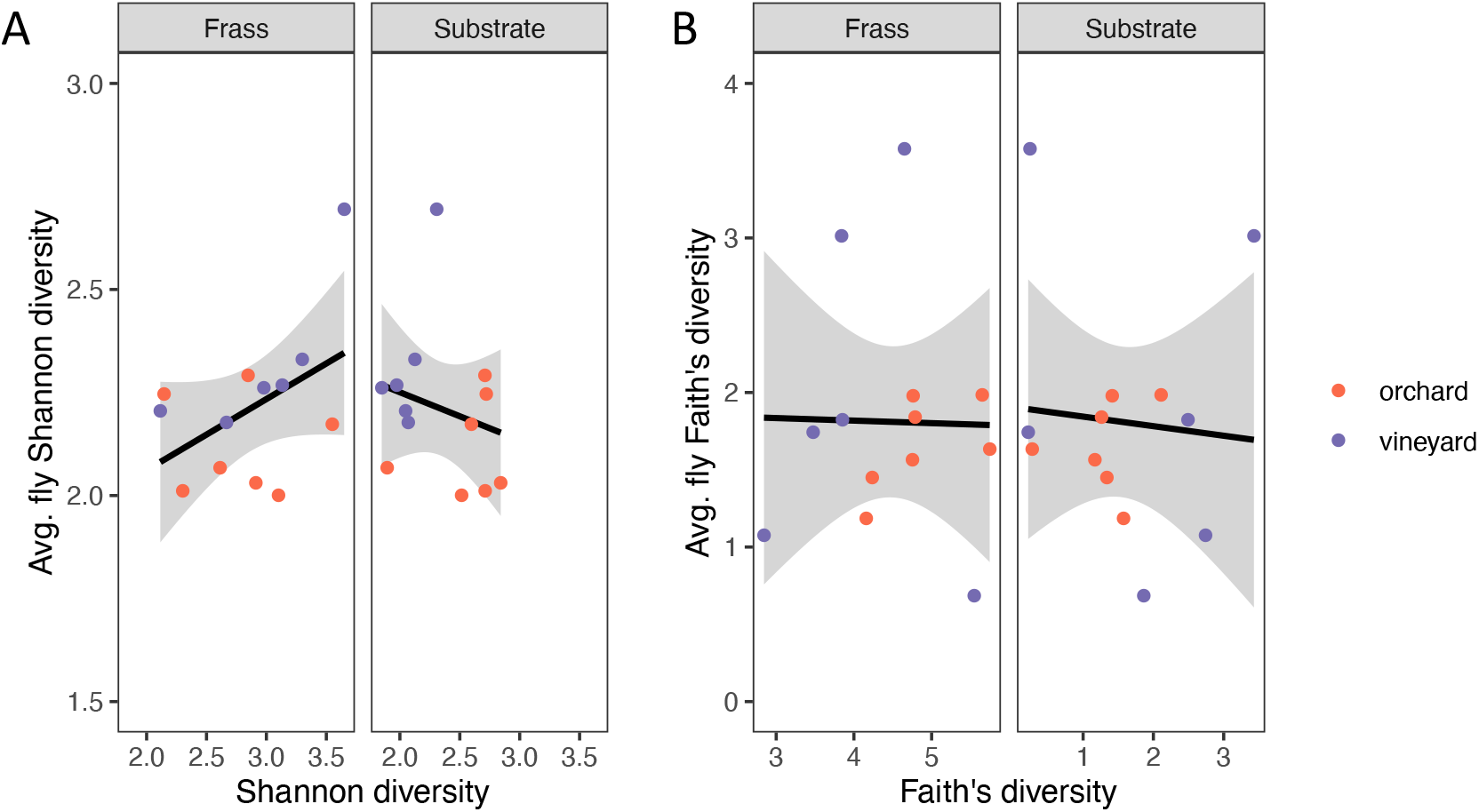
Correlations in alpha diversity between flies and either the frass or substrate. Points represent average alpha diversity measures for all flies per site, compared to the pool of frass or substrate. The model used was: alpha diversity of fly ~ alpha diversity frass/substrate + origin. A) The correlation between fly and frass Shannon diversity was not significant, though the model was marginally significant (F_2,10_ = 4.327, r^2^ = 0.36, p=0.044), driven by higher Shannon diversity in orchards. There relationship between average fly and substrate diversity differed between orchards and vineyards (t=2.542, p=0.03), though the overall model was not significant (F_2,10_ = 3.583, r^2^ = 0.309, p=0.063) B) Faith’s phylogenetic diversity was not correlated between flies and frass (F_2,10_ = 0.3319, r^2^ = −0.13, p=0.7389) or substrate (F_2,10_ = 0.3707, r^2^ = −0.12, p=0.6994).

**Supp. Fig. D1:**
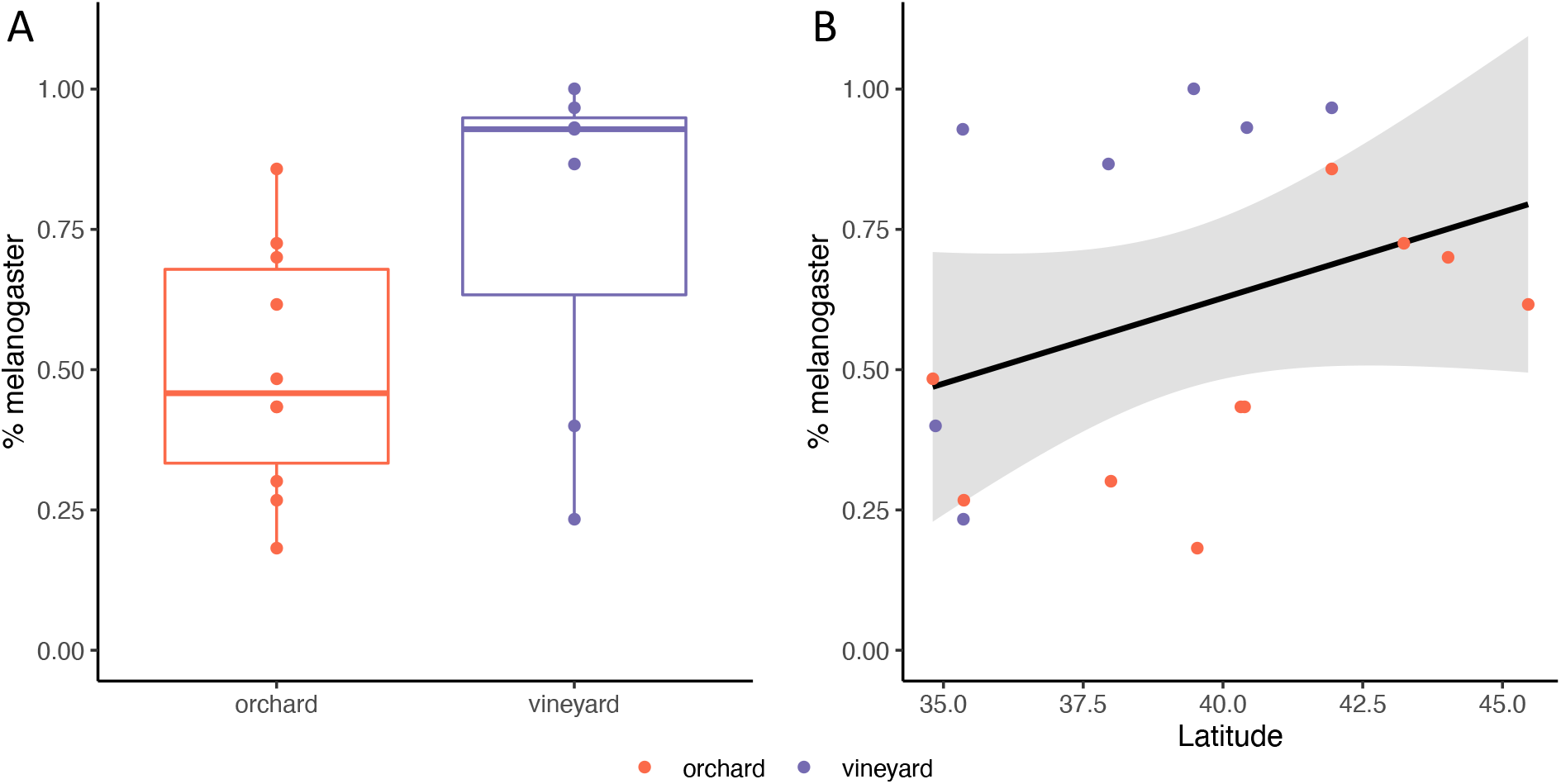
Ecological differentiation between D. melanogaster and D. simulans along the cline. Each point represents the percentage of melanogaster at all sites sampled along the cline (see Supp. Fig. M2). The data were modeled as: percent melanogaster ~ latitude + origin. A) The percent melanogaster was higher in vineyards than orchards (t = 3.537, p=0.00329). B) As latitude increases, so does the percentage of D. melanogaster (F_2,14_ = 8.237, r^2^ = 0.4762, p=0.00427).

**Supp. Fig. S1:**
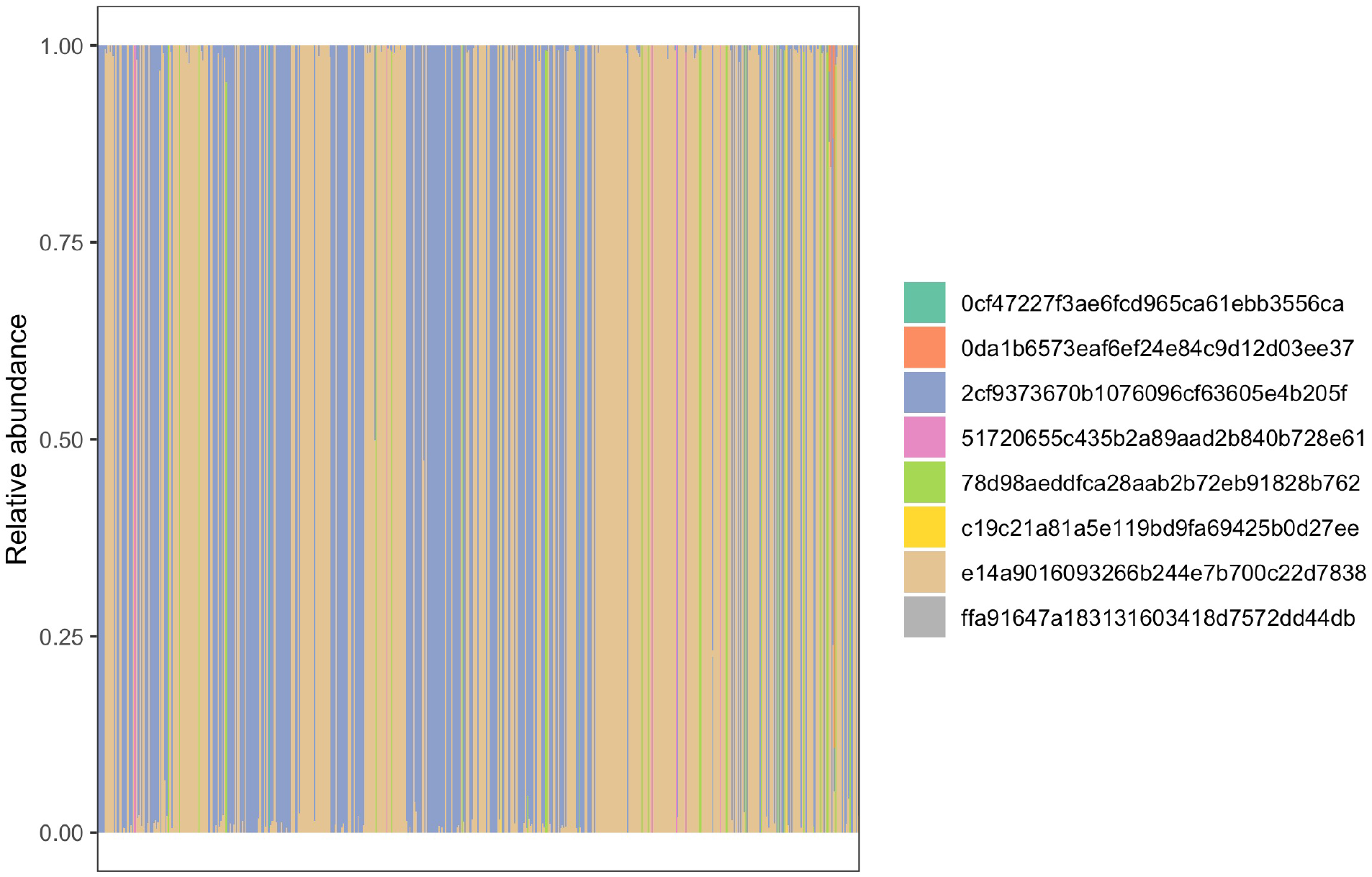
The relative abundance of COI ASVs in each individual. Each bar represents an individual, and individuals are randomly arranged along the X axis. Colors represent the relative abundance of each ASV, and most individuals had > 90% reads associated with a single ASV. 17 flies with mixed ASV were too ambiguous and were removed from subsequent analyses.

**Supp. Fig. S2:**
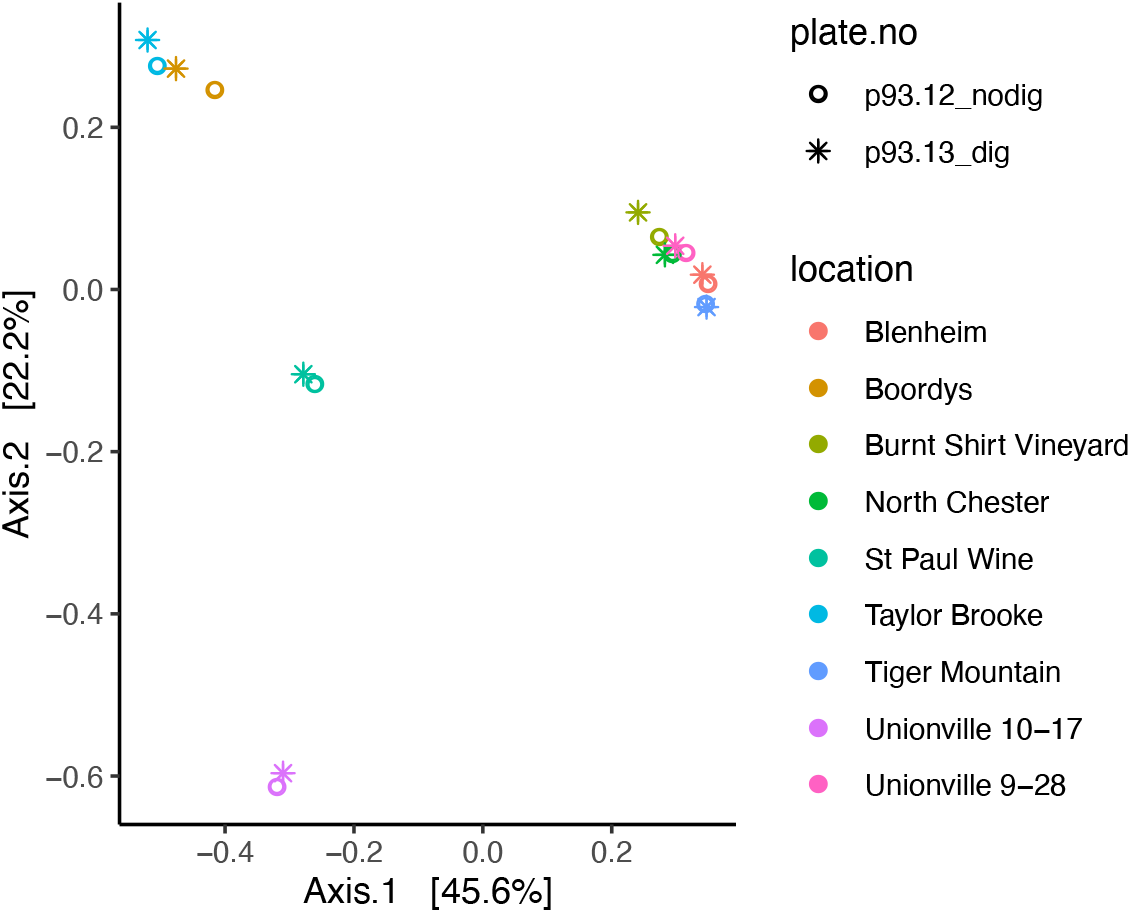
PCoA plot for the effects of BstZ17I restriction digest on substrate samples. BstZ17I digest reduces the amount of Wolbachia amplicons, but not other common bacteria in the fly microbiome. To ensure that the BstZ17I digestion did not affect the apple or grape samples, we sequenced in parallel libraries without (circles, p93.12_nodig) and with the digest (star, p93.13_dig). Points are colored by the name of the orchards or vineyard and represent PCoA based on Bray-Curtis dissimilarity. There was no significant effects of the digest on apple or grape microbiomes (PERMANOVA: F_1,16_ = 0.31, r^2^ = 0.02, p=0.91).

## References

1. McFall-Ngai M et al. 2013 Animals in a bacterial world, a new imperative for the life sciences. Proceedings of the National Academy of Sciences 110. (doi:10.1073/pnas.1218525110)

2. Koskella B, Hall LJ, Metcalf CJE. 2017 The microbiome beyond the horizon of ecological and evolutionary theory. Nat Ecol Evol 1, 1606–1615.

3. Henry LP, Bruijning M, Forsberg SKG, Ayroles JF. 2021 The microbiome extends host evolutionary potential. Nat. Commun. 12, 5141.

4. Reese AT, Dunn RR. 2018 Drivers of Microbiome Biodiversity: A Review of General Rules, Feces, and Ignorance. mBio. 9. (doi:10.1128/mbio.01294-18)

5. Gaston KJ. 2007 Latitudinal gradient in species richness. Curr. Biol. 17, R574.

6. Schemske DW, Mittelbach GG, Cornell HV, Sobel JM, Roy K. 2009 Is There a Latitudinal Gradient in the Importance of Biotic Interactions? Annu. Rev. Ecol. Evol. Syst. 40, 245–269.

7. Arnold AE, Lutzoni F. 2007 Diversity and host range of foliar fungal endophytes: are tropical leaves biodiversity hotspots? Ecology 88, 541–549.

8. Suzuki TA, Martins FM, Phifer-Rixey M, Nachman MW. 2020 The gut microbiota and Bergmann’s rule in wild house mice. Mol. Ecol. 29, 2300–2311.

9. Adrion JR, Hahn MW, Cooper BS. 2015 Revisiting classic clines in Drosophila melanogaster in the age of genomics. Trends Genet. 31, 434–444.

10. Berry A, Kreitman M. 1993 Molecular analysis of an allozyme cline: alcohol dehydrogenase in Drosophila melanogaster on the east coast of North America. Genetics 134, 869–893.

11. Paaby AB, Bergland AO, Behrman EL, Schmidt PS. 2014 A highly pleiotropic amino acid polymorphism in the Drosophila insulin receptor contributes to life-history adaptation. Evolution 68, 3395–3409.

12. Reinhardt JA, Kolaczkowski B, Jones CD, Begun DJ, Kern AD. 2014 Parallel geographic variation in Drosophila melanogaster. Genetics 197, 361–373.

13. Schmidt PS, Matzkin L, Ippolito M, Eanes WF. 2005 Geographic variation in diapause incidence, life-history traits, and climatic adaptation in Drosophila melanogaster. Evolution 59, 1721–1732.

14. Schmidt PS, Paaby AB. 2008 Reproductive diapause and life-history clines in North American populations of Drosophila melanogaster. Evolution 62, 1204–1215.

15. Douglas AE. 2019 Simple animal models for microbiome research. Nat. Rev. Microbiol. (doi:10.1038/s41579-019-0242-1)

16. Walters AW et al. 2020 The microbiota influences the Drosophila melanogaster life history strategy. Mol. Ecol. 29, 639–653.

17. Rudman SM et al. 2019 Microbiome composition shapes rapid genomic adaptation of Drosophila melanogaster. Proc. Natl. Acad. Sci. U. S. A. 116, 20025–20032.

18. Adair KL, Wilson M, Bost A, Douglas AE. 2018 Microbial community assembly in wild populations of the fruit fly Drosophila melanogaster. ISME J. 12, 959–972.

19. Simhadri RK, Fast EM, Guo R, Schultz MJ, Vaisman N, Ortiz L, Bybee J, Slatko BE, Frydman HM. 2017 The Gut Commensal Microbiome of Drosophila melanogaster Is Modified by the Endosymbiont Wolbachia. mSphere 2. (doi:10.1128/msphere.00287-17)

20. Bolyen E et al. 2019 Reproducible, interactive, scalable and extensible microbiome data science using QIIME 2. Nat. Biotechnol. 37, 852–857.

21. Callahan BJ, McMurdie PJ, Rosen MJ, Han AW, Johnson AJA, Holmes SP. 2016 DADA2: High-resolution sample inference from Illumina amplicon data. Nat. Methods 13, 581–583.

22. DeSantis TZ et al. 2006 Greengenes, a chimera-checked 16S rRNA gene database and workbench compatible with ARB. Appl. Environ. Microbiol. 72, 5069–5072.

23. McMurdie PJ, Holmes S. 2013 phyloseq: an R package for reproducible interactive analysis and graphics of microbiome census data. PLoS One 8, e61217.

24. Davis NM, Proctor DM, Holmes SP, Relman DA, Callahan BJ. 2018 Simple statistical identification and removal of contaminant sequences in marker-gene and metagenomics data. Microbiome 6, 226.

25. In press. Weather Underground. See https://www.wunderground.com/ (accessed on August 2019).

26. In press. United States Zip Codes. See https://www.unitedstateszipcodes.org/ (accessed on August 2019).

27. Oksanen J et al. 2015 Vegan community ecology package: ordination methods, diversity analysis and other functions for community and vegetation ecologists. R package ver, 2–3.

28. Bates D, Sarkar D, Bates MD, Matrix L. 2007 The lme4 package. R package version 2, 74.

29. Sloan WT, Lunn M, Woodcock S, Head IM, Nee S, Curtis TP. 2006 Quantifying the roles of immigration and chance in shaping prokaryote community structure. Environ. Microbiol. 8, 732–740.

30. Sprockett D. 2021 tyRa: Build Models for Microbiome Data.

31. Wang Y, Kapun M, Waidele L, Kuenzel S, Bergland AO, Staubach F. 2020 Common structuring principles of the Drosophila melanogaster microbiome on a continental scale and between host and substrate. Environ. Microbiol. Rep. 12, 220–228.

32. Staubach F, Baines JF, Künzel S, Bik EM, Petrov DA. 2013 Host species and environmental effects on bacterial communities associated with Drosophila in the laboratory and in the natural environment. PLoS One 8, e70749.

33. Kersters K, Lisdiyanti P, Komagata K, Swings J. 2006 The Family Acetobacteraceae: The Genera Acetobacter, Acidomonas, Asaia, Gluconacetobacter, Gluconobacter, and Kozakia. In The Prokaryotes: Volume 5: Proteobacteria: Alpha and Beta Subclasses (eds M Dworkin, S Falkow, E Rosenberg, K-H Schleifer, E Stackebrandt), pp. 163–200. New York, NY: Springer New York.

34. Shin SC, Kim S-H, You H, Kim B, Kim AC, Lee K-A, Yoon J-H, Ryu J-H, Lee W-J. 2011 Drosophila microbiome modulates host developmental and metabolic homeostasis via insulin signaling. Science 334, 670–674.

35. Huang J-H, Douglas AE. 2015 Consumption of dietary sugar by gut bacteria determines Drosophila lipid content. Biol. Lett. 11, 20150469.

36. McKenzie JA, Parsons PA. 1972 Alcohol tolerance: An ecological parameter in the relative success of Drosophila melanogaster and Drosophila simulans. Oecologia 10, 373–388.

37. David JR, Bocquet C. 1975 Similarities and differences in latitudinal adaptation of two Drosophila sibling species. Nature 257, 588–590.

38. David JR, Allemand R, Capy P, Chakir M, Gibert P, Pétavy G, Moreteau B. 2004 Comparative life histories and ecophysiology of Drosophila melanogaster and D. simulans. Genetica 120, 151–163.

39. Daisley BA, Trinder M, McDowell TW, Collins SL, Sumarah MW, Reid G. 2018 Microbiota-mediated modulation of organophosphate insecticide toxicity by species-dependent interactions with lactobacilli in a Drosophila melanogaster insect model. Appl. Environ. Microbiol. 84. (doi:10.1128/aem.02820-17)

40. Chmiel JA, Daisley BA, Burton JP, Reid G. 2019 Deleterious Effects of Neonicotinoid Pesticides on Drosophila melanogaster Immune Pathways. mBio. 10. (doi:10.1128/mbio.01395-19)

41. Soto-Yéber L, Soto-Ortiz J, Godoy P, Godoy-Herrera R. 2018 The behavior of adult Drosophila in the wild. PLoS One 13, e0209917.

42. Markow TA. 2015 The secret lives of Drosophila flies. Elife 4. (doi:10.7554/eLife.06793)

43. van der Linde K, Steck GJ, Hibbard K, Birdsley JS, Alonso LM, Houle D. 2006 FIRST RECORDS OF ZAPRIONUS INDIANUS (DIPTERA: DROSOPHILIDAE), A PEST SPECIES ON COMMERCIAL FRUITS FROM PANAMA AND THE UNITED STATES OF AMERICA. flen 89, 402–404.

44. Grainger TN, Rudman SM, Schmidt P, Levine JM. 2021 Competitive history shapes rapid evolution in a seasonal climate. Proc. Natl. Acad. Sci. U. S. A. 118. (doi:10.1073/pnas.2015772118)

45. Burns AR, Stephens WZ, Stagaman K, Wong S, Rawls JF, Guillemin K, Bohannan BJM. 2016 Contribution of neutral processes to the assembly of gut microbial communities in the zebrafish over host development. ISME J. 10, 655–664.

46. Björk JR, Díez-Vives C, Astudillo-García C, Archie EA, Montoya JM. 2019 Vertical transmission of sponge microbiota is inconsistent and unfaithful. Nat Ecol Evol 3, 1172–1183.

47. Dobson AJ et al. 2015 Host genetic determinants of microbiota-dependent nutrition revealed by genome-wide analysis of Drosophila melanogaster. Nat. Commun. 6, 7296.

48. Obadia B, Güvener ZT, Zhang V, Ceja-Navarro JA, Brodie EL, Ja WW, Ludington WB. 2017 Probabilistic Invasion Underlies Natural Gut Microbiome Stability. Curr. Biol. 27, 1999–2006.e8.

49. Pais IS, Valente RS, Sporniak M, Teixeira L. 2018 Drosophila melanogaster establishes a species-specific mutualistic interaction with stable gut-colonizing bacteria. PLoS Biol. 16, e2005710.

50. Bruijning M, Henry LP, Forsberg SKG, Metcalf CJE, Ayroles JF. 2021 Natural selection for imprecise vertical transmission in host–microbiota systems. Nature Ecology & Evolution 6, 77–87.

## SUPPLEMENTARY REFERENCES

1. Leach R, Parsons L. 2019 Barcode Splitter, version 0.18.5 [Software].

2. Bolyen E et al. 2019 Reproducible, interactive, scalable and extensible microbiome data science using QIIME 2. Nat. Biotechnol. 37, 852–857.

3. 2018 Database resources of the national center for biotechnology information. Nucleic Acids Res. 46, D8–D13.

4. McMurdie PJ, Holmes S. 2013 phyloseq: an R package for reproducible interactive analysis and graphics of microbiome census data. PLoS One 8, e61217.

5. Simhadri RK, Fast EM, Guo R, Schultz MJ, Vaisman N, Ortiz L, Bybee J, Slatko BE, Frydman HM. 2017 The Gut Commensal Microbiome of Drosophila melanogaster Is Modified by the Endosymbiont Wolbachia. mSphere 2. (doi:10.1128/msphere.00287-17)

6. DeSantis TZ et al. 2006 Greengenes, a chimera-checked 16S rRNA gene database and workbench compatible with ARB. Appl. Environ. Microbiol. 72, 5069–5072.

7. Davis NM, Proctor DM, Holmes SP, Relman DA, Callahan BJ. 2018 Simple statistical identification and removal of contaminant sequences in marker-gene and metagenomics data. Microbiome 6, 226.

8. Nunes MDS, Wengel PO-T, Kreissl M, Schlötterer C. 2010 Multiple hybridization events between Drosophila simulans and Drosophila mauritiana are supported by mtDNA introgression. Mol. Ecol. 19, 4695–4707.

